# *In situ* cryo-EM reveals how a flagellar motor transmits high torque

**DOI:** 10.64898/2026.10.14.759264

**Authors:** Jian Yue, Wangbiao Guo, Md Khalesur Rahman, Hui Xu, Md Samiul Haque, Aaron Ye, Huaxin Yu, Jun Liu, Md A. Motaleb

## Abstract

Bacterial flagellar motors are diverse rotary machines that require robust mechanical coupling to transmit torque across components with distinct architectures, yet how such coupling is achieved remains poorly understood. Here, we determine near-atomic *in situ* structures of the periplasmic flagellar motor in the Lyme disease spirochete *Borrelia burgdorferi*, revealing how an expanded rotor is mechanically coupled to the conserved rod through symmetry breaking and local remodeling of the rotor–rod interface. Surprisingly, pentameric FrbA forms a split washer-shaped brace around the rotor–rod junction, providing a second layer of mechanical reinforcement. Deletion of *frbA* destabilizes distal flagellar structures and impairs motility, particularly under increased viscous load. FrbA homologues are broadly distributed across spirochetes, suggesting a conserved mechanism for reinforcing high-torque flagellar motors. Together, these findings reveal how symmetry breaking couples an expanded rotor to a conserved driveshaft, while a molecular split washer reinforces this interface to enable robust torque transmission and motility.

## Introduction

Bacterial flagellar motors are rotary nanomachines that must generate and transmit torque while maintaining structural integrity under repeated mechanical load^1–3^. Although core components are broadly conserved, these motors exhibit remarkable diversity in architecture and mechanical output across bacterial species^4,5^. Powered by transmembrane ion flux, the motor transmits torque through the cytoplasmic C-ring and membrane-embedded MS-ring to the rod, which serves as a driveshaft connecting the rotor to the hook and filament and ultimately drives flagellar rotation^1–3^. High-torque motors have evolved enlarged rotors, increased numbers of stator complexes, and specialized accessory structures that support greater torque-generating capacity^6–8^. How these specialized motors transmit increased torque through the conserved axial machinery while maintaining mechanical integrity remains poorly understood.

Spirochetes exemplify this mechanical challenge, with flagellar motors capable of generating torques of up to ∼4,000 pN nm, compared with ∼1,300 pN nm in *Salmonella* and *Escherichia coli*^9^. In the Lyme disease spirochete *Borrelia burgdorferi*, approximately 7–11 periplasmic flagella originate near each cell pole and form overlapping ribbons around the cell cylinder (Fig. 1a–c)^10,11^. Their rotation deforms the entire cell body into propagating waves that drive translational movement through the viscous environments encountered during the tick-vertebrate enzootic cycle^12,13^. Consistent with these mechanical demands, the *B. burgdorferi* motor contains enlarged C- and MS-rings and 16 stator complexes organized around an elaborate periplasmic collar^6,14^. Our recent study showed that the expanded MS-ring in *B. burgdorferi* comprises 41 copies of FliF^15^, compared with 34 copies in many other bacteria, including *Salmonella*^16,17^ and *Vibrio*^18^. By contrast, the axial rod is assembled from five highly conserved flagellar proteins and retains a diameter similar to that in other bacterial motors^15,19^. This architectural disparity raises a fundamental mechanical question: how do structurally diverse rotors transmit torque efficiently to a conserved driveshaft while maintaining mechanical integrity?

**Fig. 1:**
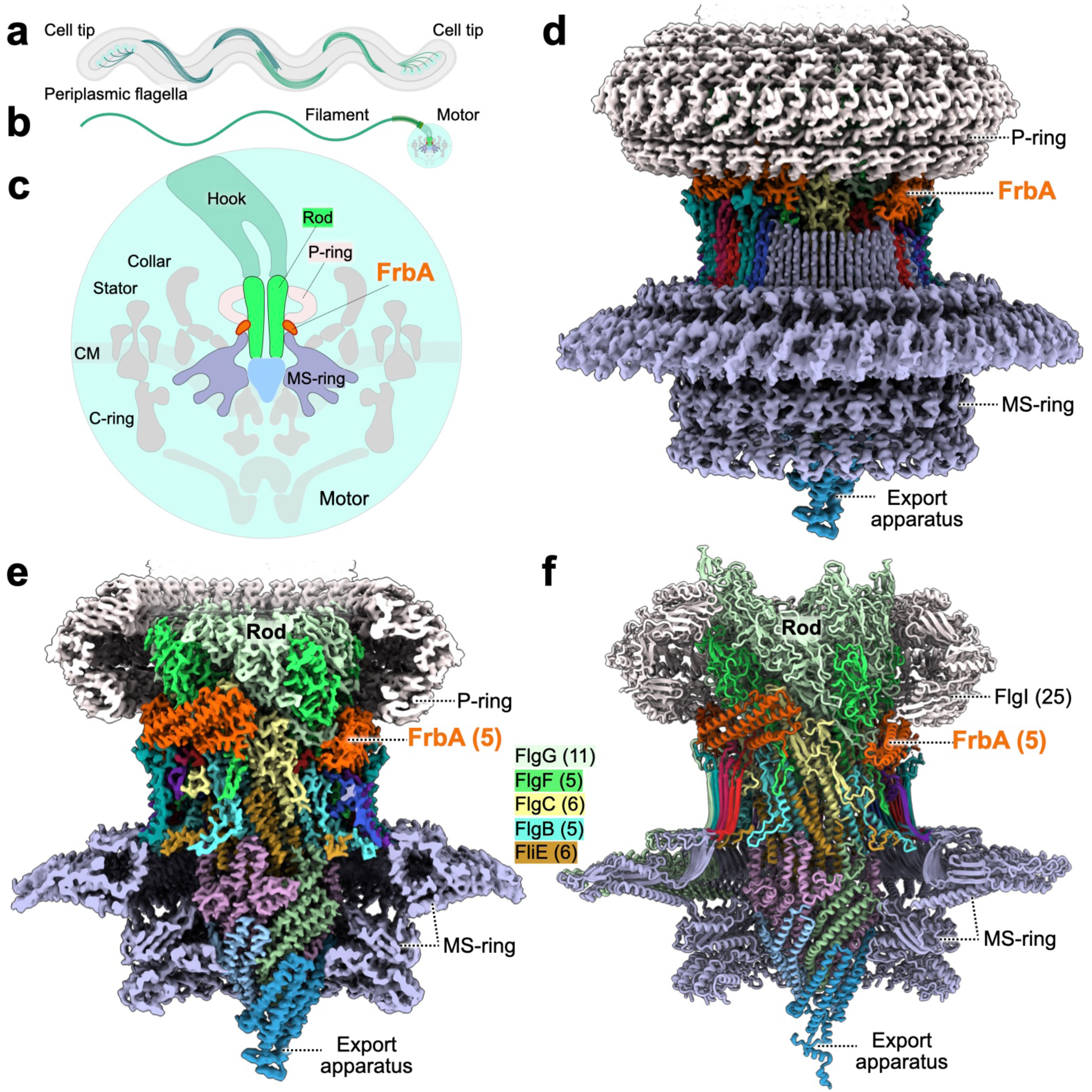
Near-atomic *in situ* architecture of the *B. burgdorferi* flagellar torque-transmission axis. **a**, Schematic of a *B. burgdorferi* cell, showing periplasmic flagella originating at both poles and extending along the cell cylinder. **b,** Schematic of an individual periplasmic flagellum and its motor. **c,** Cross-sectional schematic of the flagellar motor, showing the C-ring, MS-ring, stator complexes, collar, rod, P-ring and hook. FrbA (BB0792; orange) surrounds the rod between the MS- and P-rings. CM, cytoplasmic membrane. **d,** Composite in situ cryo-EM density map of the torque-transmission axis, comprising the MS-ring, rod, FrbA and P-ring, with the export apparatus also indicated. **e,** Central section through the density map, highlighting the axial rod and its organization relative to FrbA, the MS-ring and the P-ring. **f,** Corresponding structural model. FrbA is colored in orange, the MS-ring light purple, the P-ring light pink.

Here, we determine the near-atomic resolution structure of the torque-transmission axis in the *B. burgdorferi* flagellar motor using *in situ* single-particle cryo–electron microscopy (cryo-EM). We show that the expanded 41-subunit MS-ring undergoes extensive local remodeling to establish structurally diverse contacts with the conserved helical rod. We further identify a previously uncharacterized flagellar component, FrbA. Five FrbA molecules form a split washer-like brace around the proximal–distal rod junction, simultaneously engaging the rod and MS-ring to reinforce the torque-transmission interface. Loss of FrbA compromises axial integrity and impairs motility, while the broad conservation of FrbA across spirochetes suggests that this structural reinforcement represents a conserved adaptation of spirochetal flagellar motors. Together, our findings reveal how local remodeling and an accessory molecular brace mechanically integrate an expanded rotor with a conserved axial driveshaft, providing a structural basis for robust torque transmission.

## Results

### *In situ* cryo-EM resolves the torque-transmission axis of the *B. burgdorferi* flagellar motor

To define the architecture of the torque-transmission axis in intact motors, we used *in situ* single-particle cryo-EM to visualize periplasmic flagellar motors at the cell tips of *B. burgdorferi* (Fig. 1a–c). From 42,008 micrographs, we identified 55,867 motor particles and resolved the MS-ring–rod complex at 3.5-Å resolution, enabling atomic modeling of both assemblies (Fig. 1d,e; Extended Data Fig. 1,2). Consistent with our recent structural analysis^15^, the MS-ring comprises 41 copies of FliF. Whereas our previous reconstruction imposed C41 symmetry^15^, C1 refinement revealed pronounced local asymmetry at the MS-ring–rod junction, resolving previously obscured interactions between the MS-ring and rod.

The proximal rod comprises six FliE, five FlgB, and six FlgC subunits, whereas the distal rod contains five FlgF and FlgG subunits (Fig. 1f and 3a). Above the P-ring, axial density became progressively diffuse toward the curved hook–filament assembly, consistent with increasing conformational heterogeneity^19^. Focused refinement of the P-ring yielded a 4.4-Å reconstruction containing 25 copies of FlgI (Fig. 1d). This structure and stoichiometry differ from that of the canonical LP-ring^16,17^, which contains 26 copies each of FlgI and FlgH (Extended Data Fig. 3), revealing another architectural adaptation of the *B. burgdorferi* periplasmic flagellar motor, which lacks the L-ring^19^.

Notably, the reconstruction revealed an additional density partially encircling the rod between the MS- and P-rings that could not be assigned to any known flagellar component. This density was sufficiently well resolved for *de novo* model building with ModelAngelo^20^, which identified BB0792 as the best-matching sequence. Independent analyses using EMprot^21^ supported this assignment, and an AlphaFold3^22^-predicted model of BB0792 closely matched the cryo-EM density, including residue-level structural features (Extended Data Fig. 2a). Independently, a subtractive bioinformatics strategy designed to identify previously unrecognized flagellar proteins also identified BB0792 as a candidate (Extended Data Fig. 4). Thus, these complementary structure- and sequence-based approaches independently identified BB0792 as a previously unrecognized component of the *B. burgdorferi* flagellar motor, which we designate flagellar rod brace protein A (FrbA).

### Local remodeling couples the expanded MS-ring to the conserved flagellar driveshaft

Our near-atomic *in situ* structure revealed an extensive and highly asymmetric interface between the 41-subunit MS-ring and the rod (Fig. 2a). Although the third ring-building motif (RBM3) of FliF retains the expanded architecture of the MS-ring^15^, the β-collar undergoes pronounced local remodeling around the proximal rod (Fig. 2a–e). Two β-sheet elements deviate from the otherwise regular β-collar arrangement and bend inward toward the rod axis (Fig. 2c,d,i). Consequently, the β-collar adopts an irregular, approximately pentagonal contour spanning ∼100 Å, while inward-projecting FliF loops further constrict the upper opening to ∼54 Å (Fig. 2b,c).

**Fig. 2:**
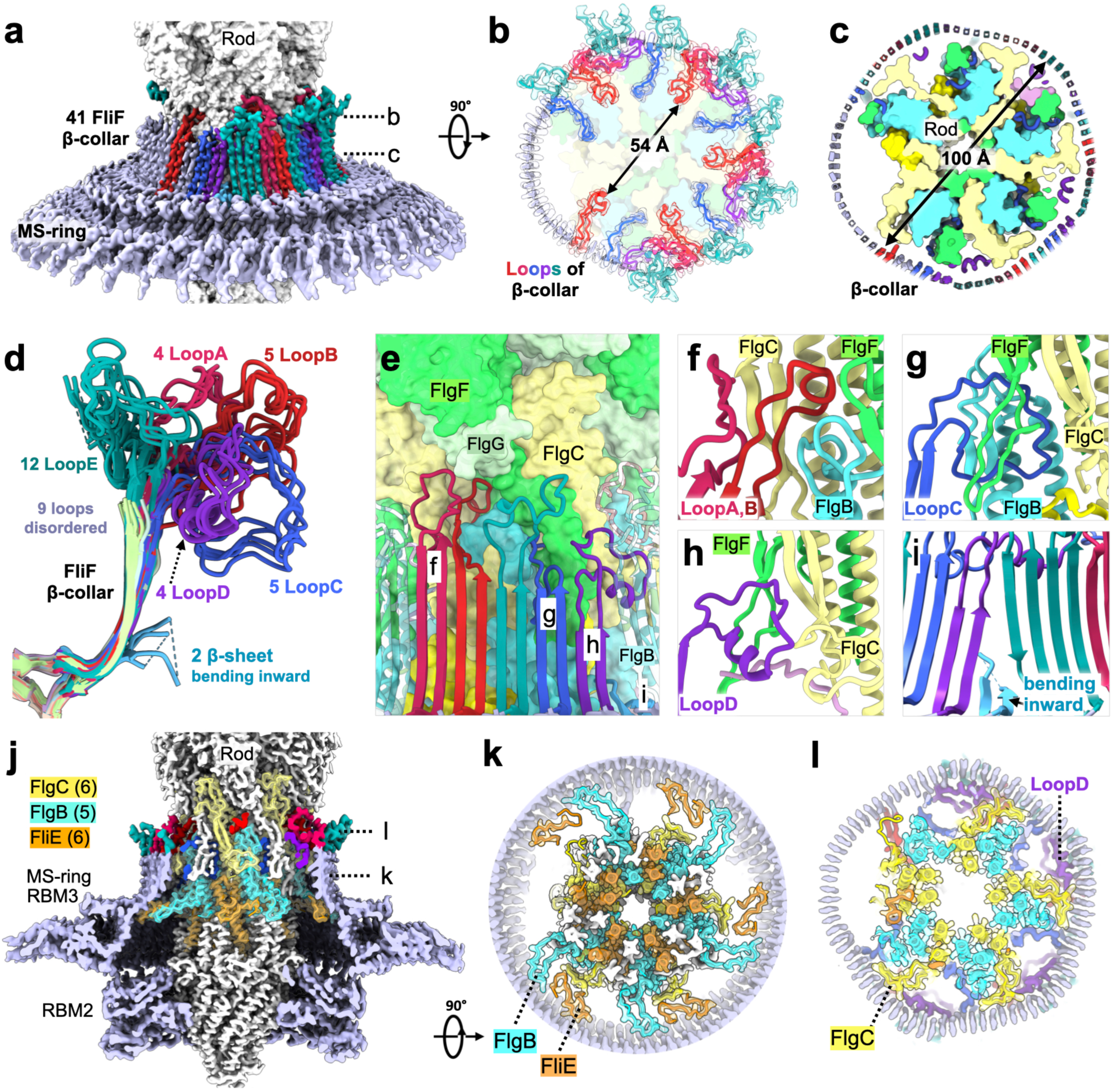
Extensive local remodeling couples the expanded MS-ring to the conserved flagellar driveshaft. **a**, Side view of the MS-ring–rod density map, highlighting the asymmetric β-collar formed by 41 FliF subunits. **b, c,** Axial sections through the upper FliF loops (**b**) and β-collar (**c**) at the levels indicated in **a**, with diameters of 54 Å and 100 Å, respectively. **d,** Superposition of FliF β-collar units, revealing conformational diversity among the extended loops, grouped into LoopA–E, and two β-sheet elements that bend inward toward the rod, whereas nine loops are disordered. **e,** Side view of the MS-ring–rod interface, with FliF elements shown as ribbons against the rod surface. **f–i** Enlarged views highlight distinct loop conformations and their local interactions with the rod. **j,** Cutaway view of the MS-ring–rod density map, showing rod elements projecting outward toward the surrounding MS-ring. RBM2 and RBM3, ring-building motif domains 2 and 3 of FliF. **k, l,** Axial sections at the levels indicated in **j**, showing contacts between the MS-ring and rod proteins FliE, FlgB and FlgC near the base (**k**) and tip (**l**) of the β-collar. Atomic models shown within the transparent cryo-EM density-map envelope.

This remodeling is accompanied by pronounced conformational diversification of a long FliF loop extending from the β-collar (residues 325–354) (Fig. 2d). Rather than adopting equivalent conformations around the 41-fold MS-ring, these loops project toward distinct features of the rod (Extended Data Movie1). Of the 41 FliF loops, 32 were sufficiently ordered for near-complete modeling, whereas nine exhibited little or no ordered density (Fig. 2b). Notably, the nine poorly ordered loops cluster within a single sector of the β-collar that lacks an associated FrbA density (Fig. 2b and 1f).

Based on their conformations and interactions, the 32 resolved loops can be grouped into five major classes, designated LoopA–E (Fig. 2d,e). Five LoopA and four LoopB elements extend into deep grooves formed primarily by FlgB, FlgC, and FlgF (Fig. 2f; Extended Data Fig. 5). Five LoopC elements penetrate inter-subunit spaces within the rod and contact both FlgB and FlgF (Fig. 2g; Extended Data Fig. 5), whereas four LoopD elements fold into the space between the MS-ring and rod, occupying local cavities as wedge-like packing motifs (Fig. 2h; Extended Data Fig. 5). These diverse interactions recur at different positions around the rod, allowing the 41-fold MS-ring to conform locally to the helical arrangement of rod subunits. By contrast, LoopE elements extend outward from the β-collar toward FrbA, establishing a peripheral connection between the MS-ring and FrbA (Fig. 2b,d).

The FliF loops therefore do not simply contact the rod surface but interdigitate with grooves, cavities, and inter-subunit spaces within the proximal rod. This coupling is reciprocal: extended elements from five FliE and five FlgB subunits project outward from the rod to contact the inner surface of the FliF β-collar, while five FlgC subunits provide additional contacts farther along the axial assembly (Figs. 2j,k,l; Extended Data Fig. 6). Together, inward-projecting FliF loops and outward-projecting rod elements create a multilayered interface spanning ∼30,700 Å^2^, approximately 3.5 times the area of the interface between the FliF RBM2 and the underlying FliPQR–FlhB export apparatus (∼8,860 Å^2^) (Fig. 2; Extended Data Fig. 6a). This symmetry-breaking architecture establishes geometric complementarity between the expanded 41-subunit MS-ring and the conserved helical driveshaft, providing a structural basis for torque transmission from the rotor to the axial rod.

### FrbA forms a pentameric mechanical brace at the proximal–distal rod junction

FrbA assembles into a pentameric, split washer-like structure that partially encircles the proximal–distal rod junction through extensive interactions with FlgC, FlgF, and FlgG (Fig. 3a,b). This assembly locally expands the diameter of the rod-associated structure to approximately 133 Å, forming an external brace around the axial driveshaft (Fig. 3c,d; Extended Data Movie 1).

**Fig. 3:**
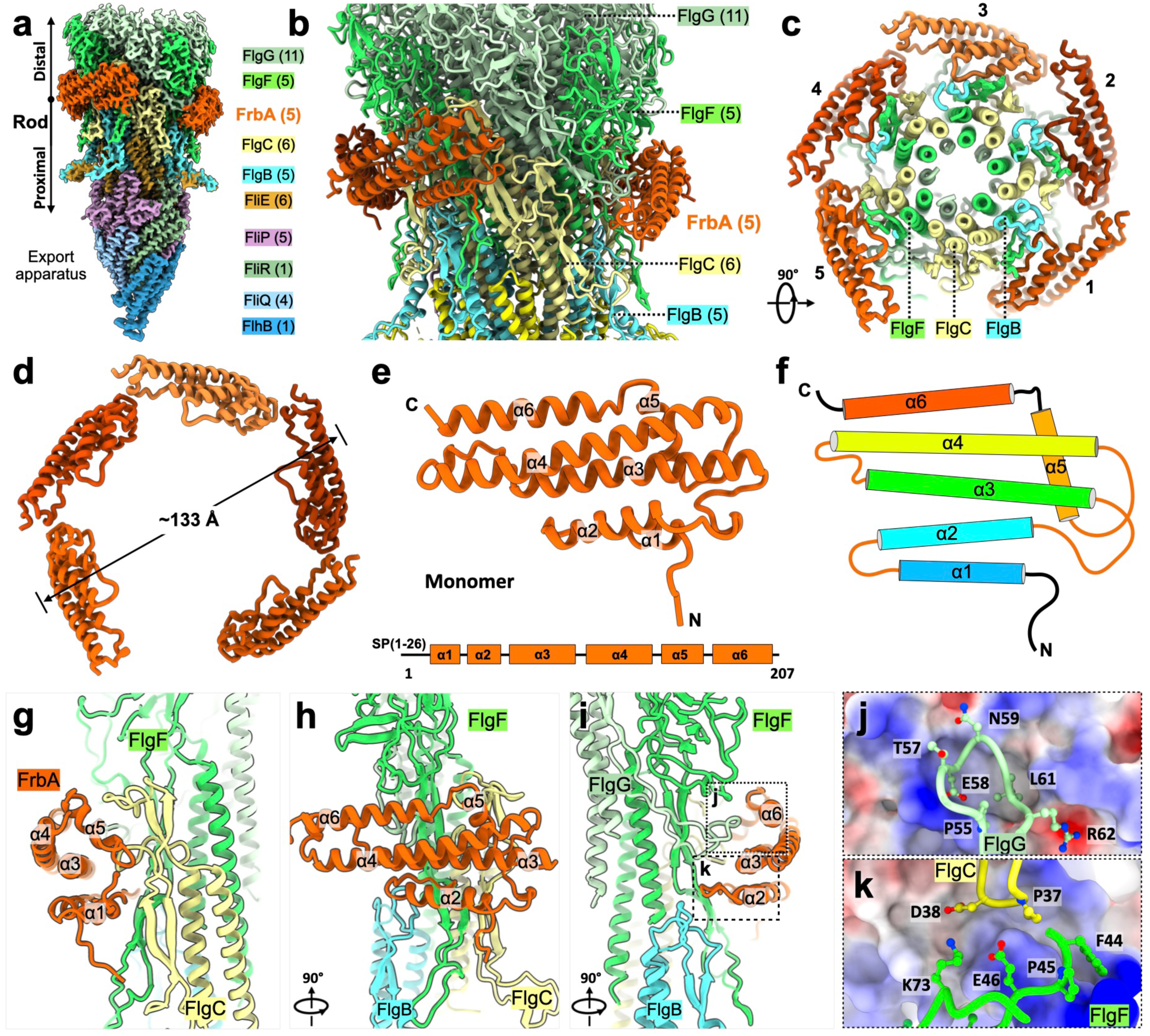
FrbA forms a pentameric mechanical brace at the proximal–distal rod junction. **a**, Density map of the rod assembly, showing FrbA (orange) at the junction between the proximal and distal rod. **b,** Enlarged view of the model, showing the FrbA assembly partially encircling the rod, with FlgB, FlgC, FlgF and FlgG indicated. **c, d,** Axial views of the five FrbA protomers assembled around the rod (**c**) and of the FrbA pentamer alone (**d**). **e, f,** Structure of FrbA monomer and its 2D topology diagram, showing its six α-helices (α1–α6) organization. SP, signal peptide. **g–i,** Views of the FrbA–rod interface related by the indicated 90° rotations, showing interactions of FrbA with FlgC, FlgF and FlgG. Boxed regions in **i** are enlarged in **j** and **k**. **j, k,** Close-up views of detail interactions between FrbA and rod residues at the interface.

Each FrbA protomer is predominantly α-helical and comprises six helices packed into an elongated bundle (Fig. 3e,f). Five protomers associate head-to-tail to form the split washer-like assembly. Rather than forming a planar ring, the protomers follow the helical organization of the underlying rod, with each engaging a corresponding set of FlgC, FlgF, and FlgG subunits. This arrangement creates five repeated interaction modules along the rod surface, with the FrbA pentamer rising helically approximately 32 Å along the rod axis (Fig. 3b).

The inner surface of the FrbA brace is formed primarily by helices α1, α5, and α6 and their connecting loops, which together constitute the principal rod-binding interface (Fig. 3g–i). This surface engages exposed regions of FlgC, FlgF, and FlgG through an extensive network of predominantly polar and electrostatic interactions (Fig. 3j,k). For example, Asp38 of FlgC, Glu46 and Lys73 of FlgF, and Glu58 and Arg62 of FlgG project toward complementary grooves on FrbA, where they are positioned to form hydrogen bonds or salt bridges. Through these multivalent interactions, each FrbA protomer simultaneously engages multiple rod components, mechanically coupling the proximal and distal rod and providing a structural basis for reinforcement of their junction.

Notably, the five FrbA protomers do not form a closed ring but leave a lateral opening approximately equivalent to the width of one additional protomer. This opening is oriented in the direction the axial assembly bends toward the periplasmic hook–filament structure (Fig. 1d, 3a–c). The split-washer architecture may therefore provide mechanical reinforcement around most of the rod circumference while preserving local flexibility along the direction of axial bending.

### FrbA braces FliF loops against the rod to reinforce the MS-ring–rod interface

Beyond its contacts with the rod, FrbA also engages FliF loops extending from the MS-ring β-collar, forming a peripheral brace around the MS-ring–rod interface (Fig. 4a–c; Extended Data Movie 1). The five FrbA protomers bury ∼8,678 Å^2^ of surface area with the rod and ∼12,583 Å^2^ with the FliF loops. On the inner side, FrbA contacts FliF Loop A–D, constraining these loops against the rod and stabilizing their rod-engaged conformations (Fig. 4d–g). On the outer side, FrbA engages the outward-projecting Loop E (Fig. 4f,g). Together, these interactions create a clamp-like arrangement in which the FliF loops interlock with the rod internally, while FrbA restrains and braces the loop network externally, thereby reinforcing the MS-ring–rod junction (Fig. 4).

**Fig. 4:**
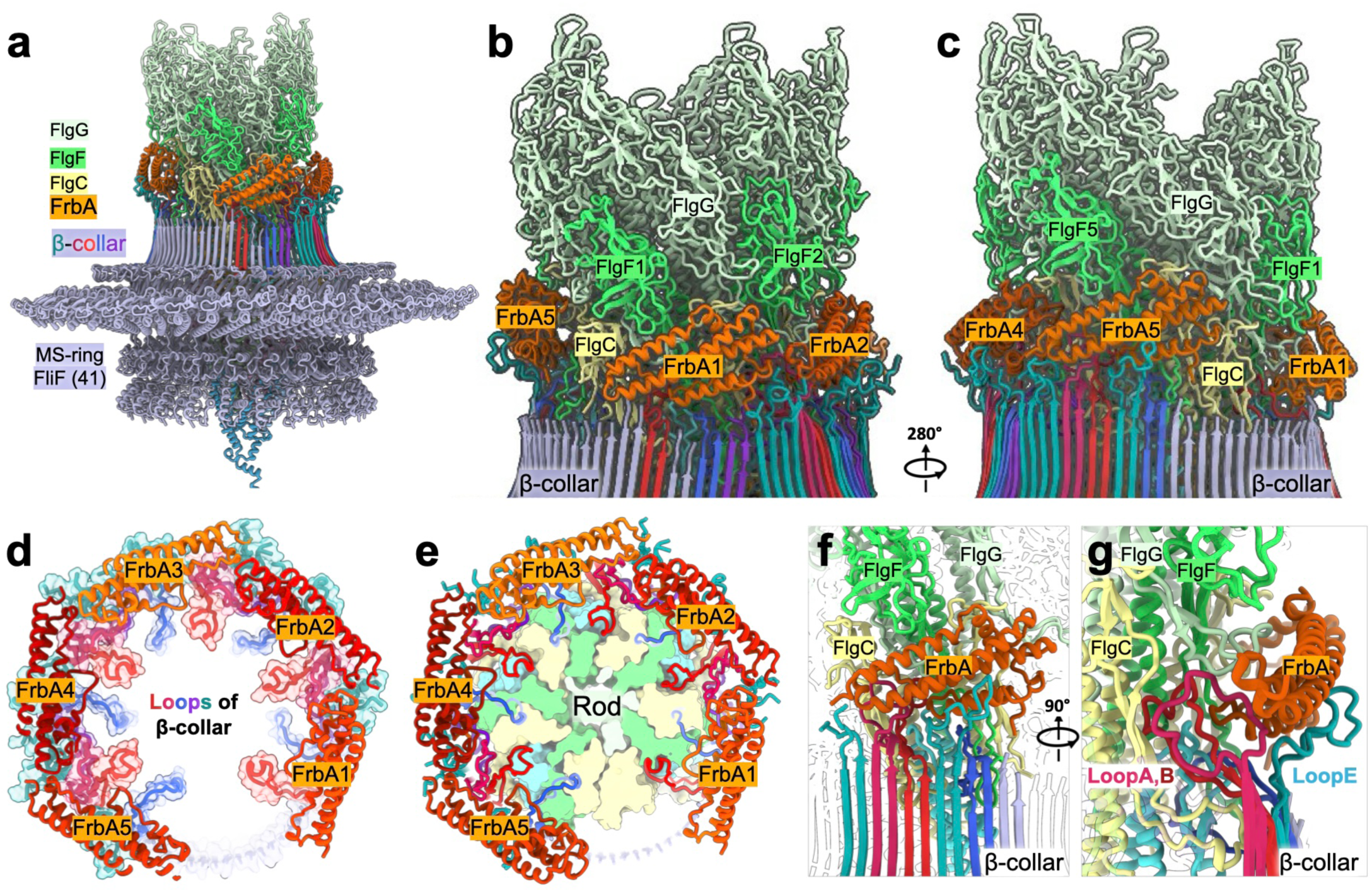
FrbA reinforces the MS-ring–rod torque-transmission interface. **a**, Overview of the MS-ring–rod model, showing the FrbA pentamer (orange) positioned above the FliF β-collar. **b, c,** Enlarged views from opposite sides of the assembly, showing FrbA interactions with both the rod and loops of FliF β-collar. Selected FrbA and FlgF protomers are numbered to indicate their positions within the assembly. **d, e,** Axial views of FrbA and the FliF loops, with the rod omitted in **d** and shown in cross-section in **e**. The FrbA pentamer partially encircles the rod junction, contacts FliF loops, and leaves a small gap in the brace. **f, g,** Enlarged views of FrbA interactions with the rod and FliF loops, highlighting how FrbA bridges the two assemblies.

The distribution of FrbA also correlates with the local ordering of the FliF loops. Most loops underlying the FrbA are well resolved, whereas the nine poorly ordered loops cluster within the opening of the FrbA pentamer (Fig. 4d,e). This spatial correlation suggests that FrbA contributes to stabilizing rod-engaged FliF conformations and helps maintain the ordered loop network around most of the MS-ring–rod interface. By simultaneously engaging the rod and the rod-contacting FliF loops, FrbA strengthens connectivity across the junction and reinforces mechanical coupling between the expanded MS-ring and the conserved flagellar driveshaft.

### FrbA contributes to flagellar axial integrity and motility

To determine the functional contribution of FrbA, we disrupted the *frbA* locus with a promoterless kanamycin Pl-Kan cassette to generate a Δ*frbA* strain. We first examined the structural consequences of FrbA loss by cryo–electron tomography (cryo-ET), which revealed frequently detached filaments from motor near the cell tips (Fig. 5a,b). Subtomogram analyses showed that 52% of Δ*frbA* motors retained an architecture comparable to wild type, whereas 34% lacked a clearly resolved hook–filament structure and 14% lacked a recognizable P-ring density (Fig. 5c–e). Genetic complementation *in cis* increased the apparently intact population to 75%, with 9% lacking a clear hook–filament structure and 16% lacking a recognizable P-ring density (Fig. 5f). Intriguingly, flagellar filaments with abnormal orientations frequently appeared in Δ*frbA* cells (Fig. 5b). Together, these structural defects indicate that FrbA contributes to the integrity and the proper orientation of periplasmic flagellar filaments.

**Fig. 5:**
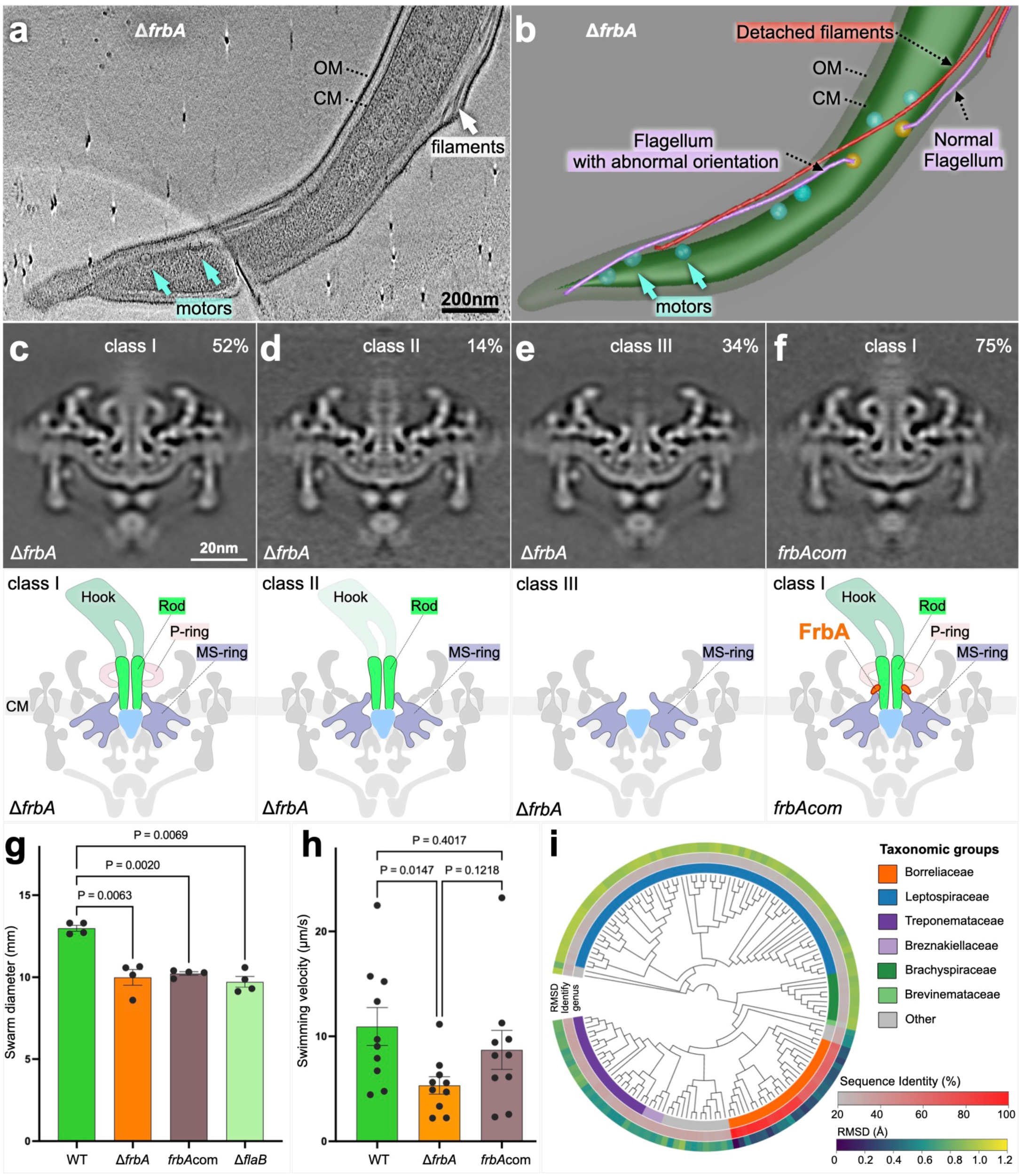
FrbA contributes to flagellar axial integrity and motility and is broadly conserved across spirochetes. **a**, Cryo-ET slice through a Δ*frbA* cell tip. Arrows indicate motors disconnected from flagellar filament. OM, outer membrane; CM, cytoplasmic membrane. **b,** Segmentation indicated the detached filaments and flagella with normal or abnormal orientations. **c–e,** Central sections through subtomogram averages and corresponding schematics of three Δ*frbA* motor classes: class I, with an apparently intact axial architecture (52%; **c**); class II, lacking recognizable P-ring density (14%; **d**); and class III, lacking clearly resolved rod–hook–filament and P-ring densities (34%; **e**). **f,** Class I subtomogram average of the complemented strain (*frbA*com), representing 75% of classified motors. g, Swarm diameters on 0.35% soft agarose. Data are mean ± s.d. from four plates. P values for the indicated comparisons were determined using paired Student’s *t*-tests. **h,** Swimming velocities of wild-type (WT), Δ*frbA* and complemented cells in 1% methylcellulose. Data are mean ± s.<u>e</u>.; 10 cells were analyzed per strain. P values for the indicated comparisons were determined using Welch’s *t*-tests. **i,** Phylogenetic distribution of FrbA homologues across 149 representative Spirochaetota taxa. The inner ring indicates taxonomic grouping, the middle ring shows percentage sequence identity relative to *B. burgdorferi* FrbA, and the outer ring shows structural RMSD relative to FrbA.

On agarose plates, Δ*frbA* cells exhibited a significant reduction in motility relative to wild type (*P* = 0.006) (Fig. 5g; Extended Data Fig. 7a). We next measured swimming velocity in methylcellulose to assess motility under increased viscous load. Under these conditions, Δ*frbA* cells showed a marked reduction in swimming velocity relative to wild type (Fig. 5h; Extended Data Fig. 7b). Thus, loss of FrbA compromises flagellar function, with a more pronounced phenotype under increased mechanical load. Genetic complementation of Δ*frbA* could partially restore structure and motility (Fig. 5g,h), likely because flagellar proteins form complex assemblies that depend on precise stoichiometric ratios to ensure structural integrity and function^23–25^.

Together, the structural and motility phenotypes support a role for FrbA in reinforcing the flagellar torque-transmission machinery. In the absence of FrbA, flagellar motor assembly is maintained, but axial structures are more fragile, and motility is impaired, consistent with reduced mechanical stability of the flagellar axial complex.

### FrbA homologues are broadly conserved across spirochetes

To examine the evolutionary distribution of FrbA, we performed sequence- and structure-based homology searches across the phylum Spirochaetota. FrbA homologues were identified in 149 species from diverse genera, including *Borreliella*, *Borrelia*, *Treponema*, *Leptospira,* and *Spirochaeta*, indicating that FrbA is broadly distributed across major spirochetal lineages (Fig. 5i; Extended Data Table 2). Despite low sequence similarity, with many homologues sharing only ∼25% identity with *B. burgdorferi* FrbA, AlphaFold3-predicted structures retained a highly similar α-helical architecture (Fig. 5i; Extended Data Fig. 8). Structural superposition yielded RMSD values below 1.2 Å, revealing strong structural conservation. Thus, these findings suggest that FrbA represents a structurally conserved component of spirochetal flagellar motors and may support a broadly conserved mechanism for reinforcing the high-torque torque-transmission machinery.

## Discussion

Efficient torque transmission requires robust mechanical coupling between the rotor and axial driveshaft while accommodating species-specific architectural adaptations. Our near-atomic *in situ* structures reveal how the expanded *B. burgdorferi* flagellar motor meets this mechanical challenge without requiring a corresponding expansion of the conserved rod. Instead, the expanded MS-ring undergoes extensive local remodeling: the FliF β-collar deviates from its otherwise regular organization, while its rod-facing loops adopt diverse conformations that interdigitate with grooves and inter-subunit spaces in the proximal rod. This asymmetric interface is further reinforced by FrbA, which forms a split washer-shaped molecular brace around the proximal–distal rod junction while engaging both the rod and MS-ring. Together with the structural defects and impaired motility caused by loss of FrbA, these findings reveal a two-layered strategy in which local remodeling accommodates the architectural mismatch between an expanded rotor and a conserved driveshaft, while an accessory molecular brace reinforces the resulting torque-transmission interface (Extended Data Movie 2).

Previous structures resolved ten rod-associated FliF peptides in *Salmonella*^16^ and fifteen FliF loops in *Vibrio*^18^, establishing that extended FliF loops directly contribute to coupling the MS-ring to the helical rod. Our *in situ* reconstruction resolves 30 FliF loops across the MS-ring–rod junction and reveals substantial conformational heterogeneity among otherwise identical FliF subunits. Individual loops adapt to distinct grooves, cavities, and inter-subunit spaces within the rod, whereas others extend toward FrbA. This structural plasticity provides a mechanism for accommodating the geometric mismatch between an expanded, rotationally symmetric MS-ring and a narrower helical driveshaft without significantly altering the conserved rod architecture. Thus, rather than propagating architectural changes throughout the axial driveshaft, the motor accommodates symmetry mismatch locally through conformational remodeling at the coupling interface.

FrbA adds a second layer of reinforcement through a distinct peripheral coupling pathway. Five FrbA protomers form a split washer-shaped brace around the proximal–distal rod junction, engaging FlgC, FlgF, and FlgG through repeated multivalent interactions while also contacting FliF loops. The resulting architecture establishes two complementary mechanical interfaces: an inner interface in which FliF interdigitates directly with the rod and an outer interface in which FrbA links the MS-ring to the rod. Such distributed connectivity may reduce reliance on any single interaction and help maintain mechanical continuity across the junction under load. The incomplete FrbA ring may also be functionally important: the brace reinforces most of the rod circumference while leaving a lateral opening oriented toward the direction of axial bending. This architecture may therefore combine mechanical reinforcement with the local flexibility required to orient the flagellar filament toward the cell center (Fig. 1a,b).

The broad distribution and strong structural conservation of FrbA homologues suggest that reinforcement of the axial machinery is a conserved feature of spirochetal motors. A protein occupying a similar position has recently been described in *Vibrio*, where FlrP associates with the FliF–rod interface but engages the axial machinery through a much more limited interaction network^18^. FrbA and FlrP share no detectable sequence or structural similarity yet occupy analogous positions at the MS-ring–rod junction, suggesting that distinct bacterial lineages independently evolved accessory components that reinforce this mechanically critical interface. Other high-torque motors may employ analogous mechanisms. *Helicobacter pylori* and *Campylobacter jejuni*, for example, possess elaborate motor-associated structures characteristic of their high-torque flagellar systems^5,7,26^. Determining whether these motors similarly expand FliF–rod interactions or deploy additional junctional components will establish how broadly reinforcement of the torque-transmission pathway accompanies the evolution of specialized flagellar motors.

Together, our findings suggest that the evolution of high-torque flagellar motors requires adaptation not only of the machinery that generates torque, but also of the structures that transmit it. By locally remodeling the MS-ring–rod interface and reinforcing it with an accessory molecular brace, the spirochetal motor accommodates an expanded rotor while preserving the conserved axial driveshaft. This architecture illustrates how rotary nanomachines can evolve specialized coupling interfaces to withstand increased mechanical loads. In spirochetes, reinforcement of this transmission axis may help sustain motility in the mechanically demanding environments encountered during dissemination and host colonization^27–29^.

## Materials and Methods

### Bacterial strains and growth conditions

The high-passage *B. burgdorferi* strain B31A (wild type and Δ*frbA*) was grown in BSK-II liquid medium supplemented with 6% rabbit serum or on semisolid agar plates at 35°C in 3% carbon dioxide.

### Construction of the *frbA* mutant and complemented strains

Construction of the FrbA/BB0792 inactivation plasmids, electroporation, and plating conditions were described previously^30,31^. Briefly, a 236-bp upstream and a 372-bp downstream sequence, along with the bb0792 ORF (624 bp), were PCR-amplified using Q5® High Fidelity DNA Polymerase with primers (5′–3′) PF BB0792-Pl-Kan (CGAAACAGACCCAAACATAGCTTATGAAAGAATAC) and PR-BB0792-Pl-Kan (CTCTACAAGATGTTCAATGCACCATTATCTTATAGG). The PCR product was then ligated into the pGEM-T Easy vector (Promega), yielding pGEM-T Easy::bb0792. The Pl-Kan cassette^30^ was PCR-amplified from PflgB-Kan^12^ using primers (5′–3′) PF-Pl-Kan-XbaI (GATGATGAATCTAGATAGTTAAAAGCAATTTTAAATG) and PR-Pl-Kan-XbaI (CTATTAATTTCTAGATTAGAAAAACTCATCGAGCATC). The XbaI-restricted Pl-Kan DNA was then inserted into the XbaI sites within BB0792 (located 380 and 1232 base pairs from the bb0792 ATG start codon) of the pGEM-T Easy::BB0792 vector, yielding pGEM-T Easy::BB0792-Pl-Kan. This insertion mutation was in-frame. The transcriptional orientation of Pl-Kan was verified by a second PCR using primers (5′–3′) PF outside-BB0792-(CATTTGCATATCTTTATGCTGCAGACCGACACG) and PR outside-BB0792 (GGAGCTTATTGTGGGCACATTGATTGCATGGAAG). BB0792-Pl-Kan was digested with the NotI restriction enzyme and separated from the pGEM-T Easy::BB0792-Pl-Kan vector, and the linear DNA was prepared for electroporation into competent B31-A cells. Transformants were selected on 200 μg/ml kanamycin. Kanamycin-resistant transformants were isolated, and the insertion of Pl-Kan was confirmed by PCR.

The Δ*frbA*/*bb0792* mutant strain was complemented in cis by chromosomal integration using the pXLF14301 suicide vector, as described previously^32,33^. In brief, the native promoter regions of the *bb0793* (Pbb0793) and *bb0792* genes were separately PCR-amplified from WT *B. burgdorferi* strain B31-A genomic DNA using primer pairs PF PBB0793 SpeI (GTCTAGAACTAGTCATTGTCAGAATAGGCATTGCTAGG) and P1R Pbb0793 (CATTTTTGAAAATATGCATAAATATTTTTTCTTCCTTTC), and P2F Pbb0793 (GAAAGGAAGAAAAAATATTTATGCATATTTTCAAAAATG) and PR Pbb0792NotI (CTCGGGTAGCGGCCGCTTAATAATAGTTTGGCCCTTGTTC). These fragments were fused by overlapping PCR to generate Pbb0793-*bb0792*, which was inserted into the derivative pXLF14301::PflgB-Strep^34–36^ using SpeI and NotI restriction sites, yielding pXLFbb0792. The plasmid was then electroporated into the Δ*bb0792* mutant cells, followed by selection with both kanamycin and streptomycin. Resistant clones were analyzed by PCR to verify the integration of Pbb0793-*bb0792*-PflgB-Strep into the intergenic region between *bb0445* and *bb0446*.

### Motility assays

Swarm plate assays were performed as described previously^31,37,38^. Approximately 1 × 10^9^ cells in a 5-μl volume were spotted onto 0.35% soft agarose plates containing BSK medium. Plates were incubated for five days at 35°C in a humidified incubator with 2.5% CO₂. The aflagellated Δ*flaB* was used as a control^37^. Swarm diameters were measured in millimeters from four plates and are reported as the mean ± standard deviation. Wild-type and mutant swarm diameters were compared using a paired Student’s t-test, with P ≤ 0.05 considered significant.

### Microscopy and computer-assisted motion analysis

Exponentially growing *B. burgdorferi* cells (4–6 × 10⁶ cells/ml) were examined using a Zeiss Axio Imager M1 dark-field microscope equipped with an AxioCam digital camera. Cells were mixed with 0.5% or 1% 400-mesh methylcellulose (Sigma-Aldrich) and video-recorded at room temperature (24°C). Swimming velocities were analyzed using AxioVision software as described previously^39^. At least 10 cells per strain were analyzed, and velocities are reported as the mean ± standard error in μm/s. Wild-type and mutant velocities were compared using Welch’s t-test in GraphPad 4.0, with P ≤ 0.05 considered significant.

### Frozen-hydrated sample preparation for cryo-EM

Frozen-hydrated specimens were prepared using established protocols^15,40,41^. Briefly, *B. burgdorferi* cultures were subjected to centrifugation at 5,000 g for 5 min. The resulting pellet was resuspended in 1.0 ml of PBS, centrifuged again and resuspended in ∼50 μl of PBS. For cryo-ET and cryo-EM, the cell suspension was mixed with or without 10 nm colloidal gold fiducial markers. A 5-μl aliquot of the cell suspension (OD600 ≈ 0.2) was applied to freshly glow-discharged holey carbon grids (Quantifoil) or lacey carbon grids (Ted Pella). Grids were blotted for 6–8 s using filter paper and rapidly plunge-frozen in liquid ethane using a GP2 plunger (Leica).

### *In situ* single-particle cryo-EM data collection

Data were collected on a Titan Krios electron microscope (Thermo Fisher Scientific) operated at 300 kV and equipped with a K3 direct electron detector and a GIF energy filter (Gatan). Movies were recorded in super-resolution mode at a nominal magnification of ×81,000, corresponding to a calibrated pixel size of 0.534 Å. Automated data acquisition was performed in SerialEM^42^ using a customized multishot script^41^ to target bacterial cells directly on cryo-EM grids. Each movie was recorded with a total electron dose of 70 e⁻/Å² fractionated over 40 frames, with a defocus range of −1.5 to −2.2 μm. Full data collection parameters are summarized in Extended Data Table 1.

### Flagellar particle selection and structure determination

All image processing and structural analyses were performed in CryoSPARC^43^, following the workflow summarized in Extended Data Fig. 1. To establish an automated particle-picking model, ∼1,000 flagellar motors were manually picked and used to train a neural-network model in Topaz^44^. The model was iteratively refined and applied to the full dataset. In total, 42,008 micrographs yielded 152,861 particles after Topaz-based picking. Following two-dimensional (2D) classification, 83,234 particles displaying clear flagellar motor features, including top and side views, were retained. Particles were extracted with a box size of 448 pixels at a pixel size of 1.43 Å.

An initial 3D model was generated by ab initio reconstruction with C16 symmetry. Heterogeneous refinement with C16 symmetry combined with 3D classification were then used to further classify the particles. Focused refinement of the basal-body and collar region improved alignment of intact motors, yielding a reconstruction from 76,561 particles at an overall resolution of ∼6 Å.

For high-resolution analysis of the motor axis, including the MS-ring and rod, signals corresponding to the stator and peripheral cage/collar regions were subtracted. The resulting particles were subjected to C16 symmetry expansion, followed by focused classification of the MS-ring without alignment under C1 symmetry. This analysis revealed that the RBM3 domains of FliF form a 41-fold symmetric S-ring. Subsequent focused refinement of the RBM3 region with C41 symmetry yielded a reconstruction at an overall resolution of 3.13 Å.

To resolve the symmetry mismatch between the MS-ring and the central rod, particles were further subjected to C41 symmetry expansion, followed by focused 3D classification of the central rod without imposing symmetry. Classes exhibiting well-defined rod density were selected, resulting in a final 55,867 particles. Focused local refinement under C1 symmetry, combined with local CTF refinement, further improved the reconstruction to an overall resolution of 3.50 Å.

For reconstruction of the P-ring, signals corresponding to the central rod and MS-ring were subtracted from the particles. Focused 3D classification under C1 symmetry revealed a 25-fold rotational organization of the P-ring. Subsequent local refinement with C25 symmetry yielded a final P-ring reconstruction at an overall resolution of 4.29 Å.

Local resolution was estimated in CryoSPARC^43^ using the gold-standard Fourier shell correlation (FSC) threshold of 0.143, as summarized in Extended Data Fig. 1. Final maps were post-processed with EMReady^45^ to facilitate model building and visualization. Representative model-to-map fits are shown in Extended Data Fig. 2.

### Identification of FrbA/BB0792

The EMReady-enhanced density map was imported into RELION5^46^ for ModelAngelo analysis^20^ using the complete proteome of *Borrelia burgdorferi* strain B31A (NCBI taxonomy ID: 224326) as the sequence database. The sequence assigned to the de novo model was evaluated using profile hidden Markov model (HMM) and BLAST searches, both of which identified BB0792 as the best-matching protein. Independent analysis with EMprot^21^ produced a consistent assignment. The AlphaFold3-predicted structure of BB0792 could also be fitted into the density. Manual inspection confirmed agreement between the experimental density and modeled backbone and side-chain features, supporting the identification of BB0792. Independently, a subtractive bioinformatics strategy designed to identify previously unrecognized flagellar proteins also identified BB0792 as a candidate (Extended Data Fig. 4)

### Model building and refinement

Initial atomic models were generated with AlphaFold3^22^ and fitted as rigid bodies into the cryo-EM density maps using UCSF ChimeraX^47^. The fitted models underwent molecular dynamics-based flexible fitting with Namdinator^48^ and ISOLDE^49^, followed by iterative manual adjustment and rebuilding in Coot^50^. Real-space refinement was performed in PHENIX^51^ with appropriate geometry restraints. Final models were evaluated using MolProbity^52^. Structure refinement and validation statistics are provided in Extended Data Table 1.

### Cryo-ET data collection and analyses

Frozen-hydrated specimens were imaged at -170°C using a 300 kV Krios electron microscope (Thermo Fisher Scientific) equipped with a field emission gun, Volta phase plate, and a post-GIF direct electron detector (Gatan) at a magnification of ×42,000 and a pixel size of 2.148 Å. Tilt series were collected from −48° to 48° with 3° increments using SerialEM^42^ and FastTomo^53^. The total dose of the tilt series was ∼70 e⁻/Å² at a defocus of −4.8 µm.

Images were first motion-corrected using MotionCorr2^54^ and subsequently stacked by IMOD^55^. Tilt series were aligned using fiducial markers or fiducial-free alignment by IMOD^55^. Gctf^56^ was employed to determine the defocus of each tilt image in the aligned stacks, and the “ctfphaseflip” function in IMOD^55^ was used for CTF correction of the tilt images. Tomograms were generated using Tomo3D^57^ by either simultaneous iterative reconstruction technique (SIRT) or weighted back-projection (WBP). SIRT reconstructions were utilized for particle picking of flagellar motors due to their higher contrast, while tomograms reconstructed by WBP retained high-resolution details for *in situ* structure determination of the flagellar motor through subtomogram averaging.

### Subtomogram averaging and corresponding analysis

Flagellar motors were manually picked from the tomographic reconstructions. 3D alignment and classification were performed with the protomo package as described previously^6,10,58^. Motors were initially aligned in 8× binned tomograms using their estimated orientations. The center coordinates of the C-ring and collar provided two of the three Euler angles. Subtomograms were then extracted from unbinned tomograms using the refined coordinates and binned 4× for alignment. Multi-reference alignment and 3D classification were performed as described previously^6,10,58^. This process was repeated with 2× binned subtomograms to improve structural detail.

### FrbA conservation analysis in Spirochaetota

A set of 149 Spirochaetota genomes was selected^59^, as summarized in Extended Data Table 2. Candidate FrbA/BB0792 homologues were identified using sequence similarity searches and profile-HMM searches. Local sequence alignments were performed using Biopython, and a profile HMM constructed from MAFFT-aligned BB0792 homologues was used to search the corresponding proteomes with HMMER v3.4^60^. Structure-based searches using Foldseek^61^ provided additional evidence supporting homology. Together, these analyses identified candidate homologues in all 149 genomes.

Structures of the candidate homologues were predicted using AlphaFold 3^22^. For each sequence, the model with the highest ranking score was selected and aligned to the AlphaFold 3-predicted BB0792 reference structure using MatchMaker in UCSF ChimeraX^62^. Structural comparisons were restricted to Cα atoms of residues with pLDDT scores ≥70 in both models. Core root-mean-square deviation (RMSD) was calculated. The phylogenetic tree was generated by the GTDB v232^63^ bac120 reference tree to the corresponding 149 tips. Sequence identity relative to BB0792 and core RMSD were visualized as annotation rings using iTOL^64^.

## Data availability

The atomic coordinates and corresponding density maps have been deposited in the Protein Data Bank (PDB) and the Electron Microscopy Data Bank (EMDB). FrbA–MS-ring–rod structure: PDB 39BU and EMD-79298; FrbA–MS-ring–rod structure together with the P-ring: PDB 39BV and EMD-79299; P-ring (25 FlgI) structure: PDB 39BW and EMD-79300.

## Acknowledgments

We thank Jennifer Aronson for critical reading and editing of the manuscript and Jianfeng Lin and Kangkang Song for assistance with cryo-EM data collection. We thank Beile Gao for assistance with the FrbA conservation analysis.

## Author Contributions Statement

J.Y., J.L., and M.A.M. conceived and designed the study. J.Y., W.G., and J.L. processed the cryo-EM data, determined the structures, and performed model building, structural analysis, and visualization. J.Y., W.G., M.K.R., and H.Y. prepared the cryo-EM samples, collected the data, and data analysis. M.K.R., H.X., M.S.H., and M.A.M. performed the genetic experiments and motility assays. A.Y. and J.Y. performed tomogram segmentation and movie animation. J.Y., J.L., and M.A.M. wrote the original manuscript. All authors reviewed and edited the manuscript. J.L. and M.A.M. supervised the study and acquired funding.

## Supplemental Information

Extended Data Table 1

Extended Data Table 2

Extended Data Movie 1

Extended Data Movie 2

**Extended Data Fig. 1.**
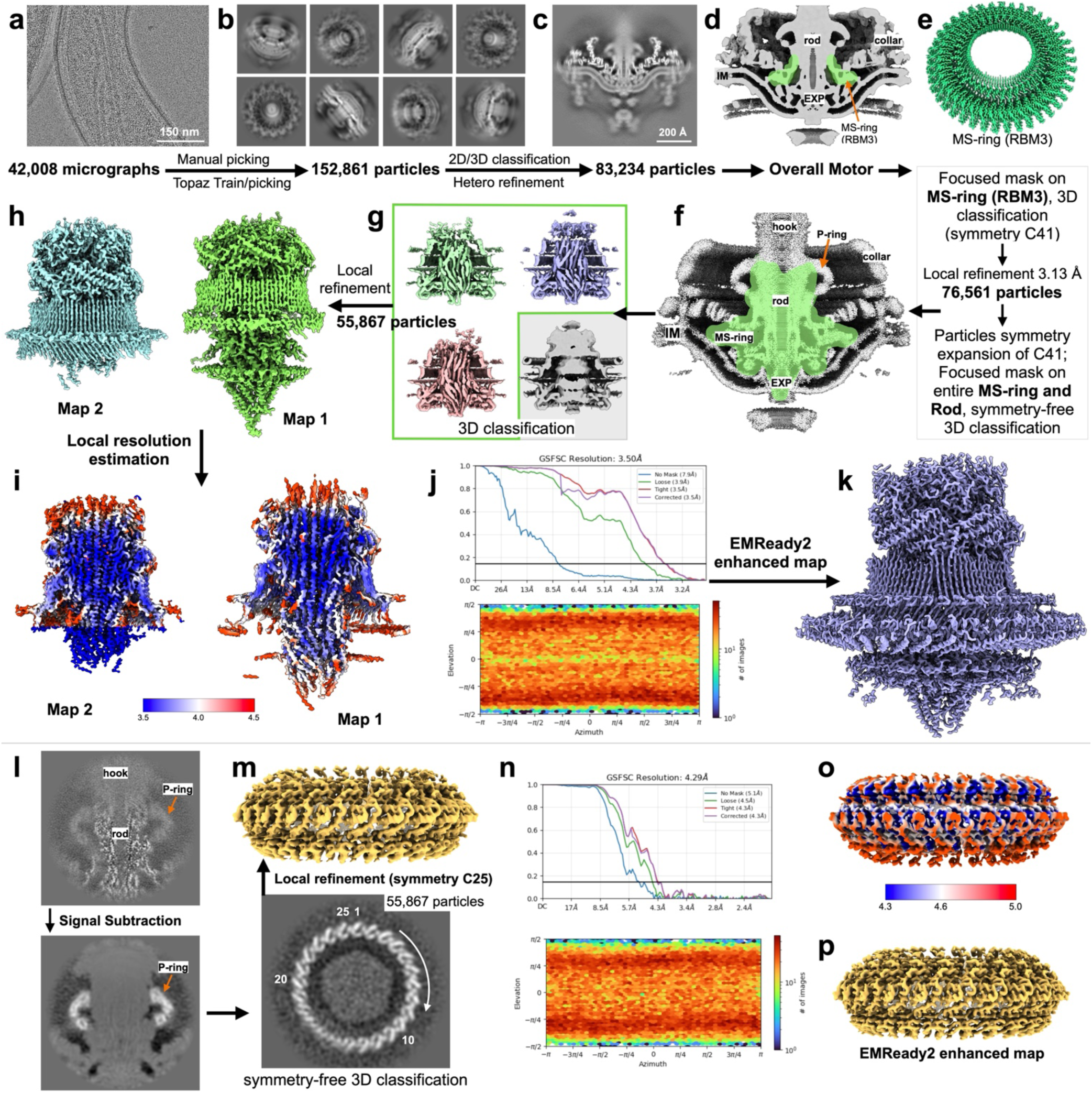
: Cryo-EM processing workflow and validation of the MS-ring–rod and P-ring reconstructions. **a**, Representative cryo-EM micrograph of *B. burgdorferi*. **b**, Representative 2D class averages. Manual picking and Topaz-assisted picking from 42,008 micrographs yielded 152,861 particles. **c**, Center slice of the motor after 2D/3D classification and heterogeneous refinement, which retained 83,234 particles. **d**, Cutaway view of the overall motor reconstruction, with the MS-ring RBM3 region highlighted in green. **e**, Focused classification and local refinement of RBM3 with C41 symmetry yielded a 3.13 Å reconstruction from 76,561 particles. **f**, Mask encompassing the MS-ring and rod for symmetry-free 3D classification after C41 symmetry expansion. **g**, Classification results, with selected classes outlined in green. **h**, Locally refined MS-ring–rod maps, designated Map 1 and Map 2; 55,867 particles were retained for local refinement. **i**, Maps colored by local resolution according to the scale (Å). **j**, Gold-standard Fourier shell correlation (FSC) curves and particle orientation distribution for the MS-ring–rod reconstruction. **k**, MS-ring–rod map enhanced using EMReady2. **l**, P-ring isolation by signal subtraction followed by symmetry-free 3D classification. **m**, P-ring reconstruction obtained by local refinement with C25 symmetry using 55,867 particles. **n**, FSC curves and particle orientation distribution for the P-ring reconstruction. **o**, P-ring map colored by local resolution (Å). **p**, P-ring map enhanced using EMReady2. IM, inner membrane; EXP, export apparatus; RBM3, ring-building motif 3.

**Extended Data Fig. 2:**
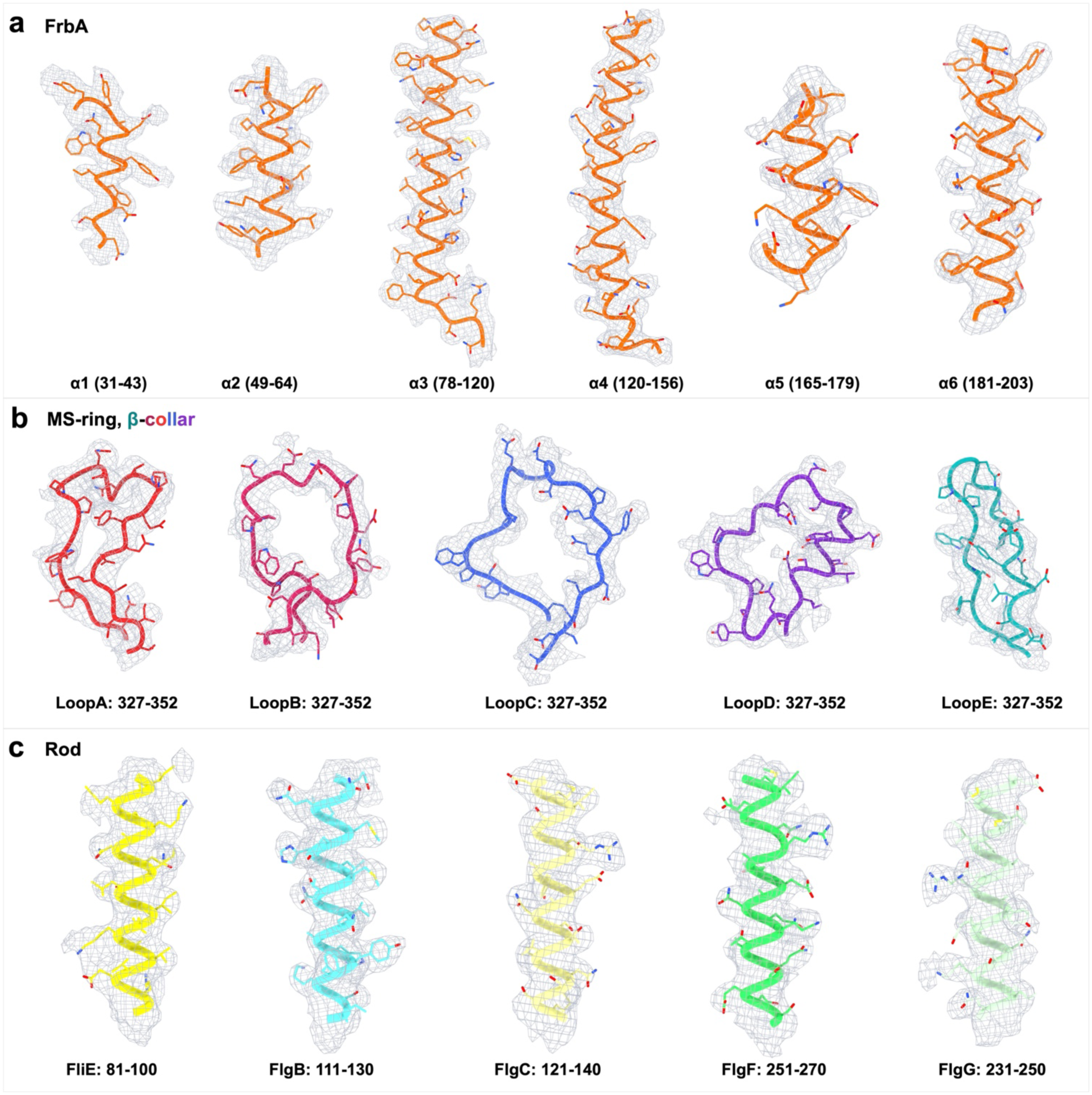
Representative cryo-EM density map-atomic model fitting. **a**, Density surrounding the six α-helical segments of FrbA (orange). **b**, Density supporting five distinct conformations of the FliF β-collar loop (residues 327–352), designated LoopA–LoopE. **c**, Representative density surrounding residues 81–100 of FliE, FlgB, FlgC, FlgF and FlgG, as labelled. Atomic models are shown as ribbons with side chains in stick representation, and cryo-EM densities are displayed as grey mesh.

**Extended Data Fig. 3:**
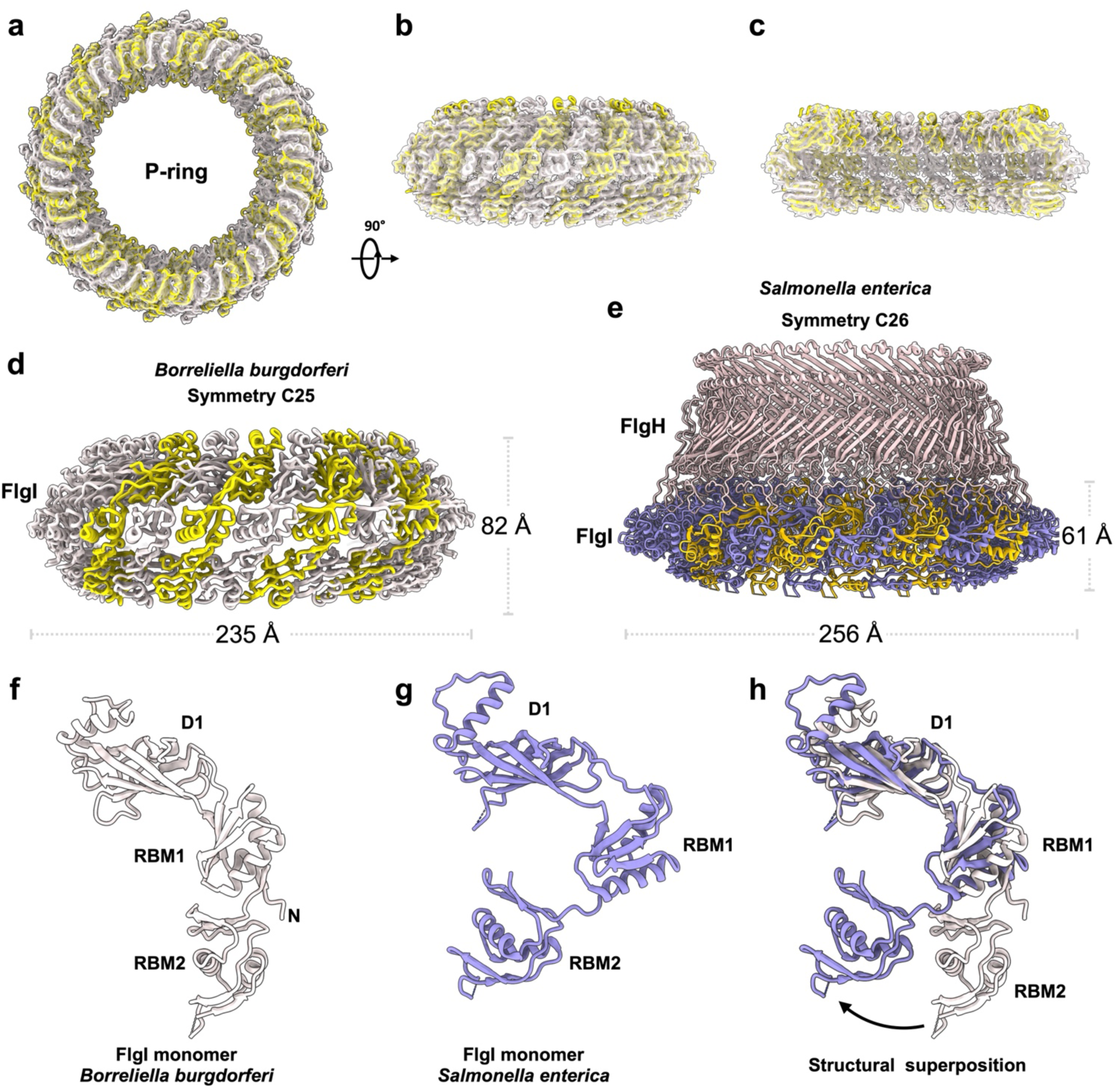
Structure of the *B. burgdorferi* P-ring and structural comparison of FlgI. **a**–**c**, Top (**a**), side (**b**) and cutaway side (**c**) views of the P-ring model fitted into the transparent cryo-EM density map. **d**, Side view of the *B. burgdorferi* P-ring, comprising 25 FlgI subunits arranged with C25 symmetry, with an outer diameter of approximately 235 Å and a height of 82 Å. **e**, Side view of the *Salmonella enterica* LP-ring complex with C26 symmetry (PDB: 7CBL). The P-ring has an outer diameter of approximately 256 Å and a height of 61 Å. Compared with the *S. enterica* P-ring, the *B. burgdorferi* P-ring is narrower and more elongated along the ring axis. **f**, **g**, FlgI monomers from *B. burgdorferi* and *S. enterica*. Domain D1 and ring-building motifs RBM1 and RBM2 are labelled. **h**, Structural superposition of the monomers shown in **f** and **g**, revealing similar individual domain structures but differences in their relative arrangement, consistent with the greater axial extension of the *B. burgdorferi* P-ring.

**Extended Data Fig. 4:**
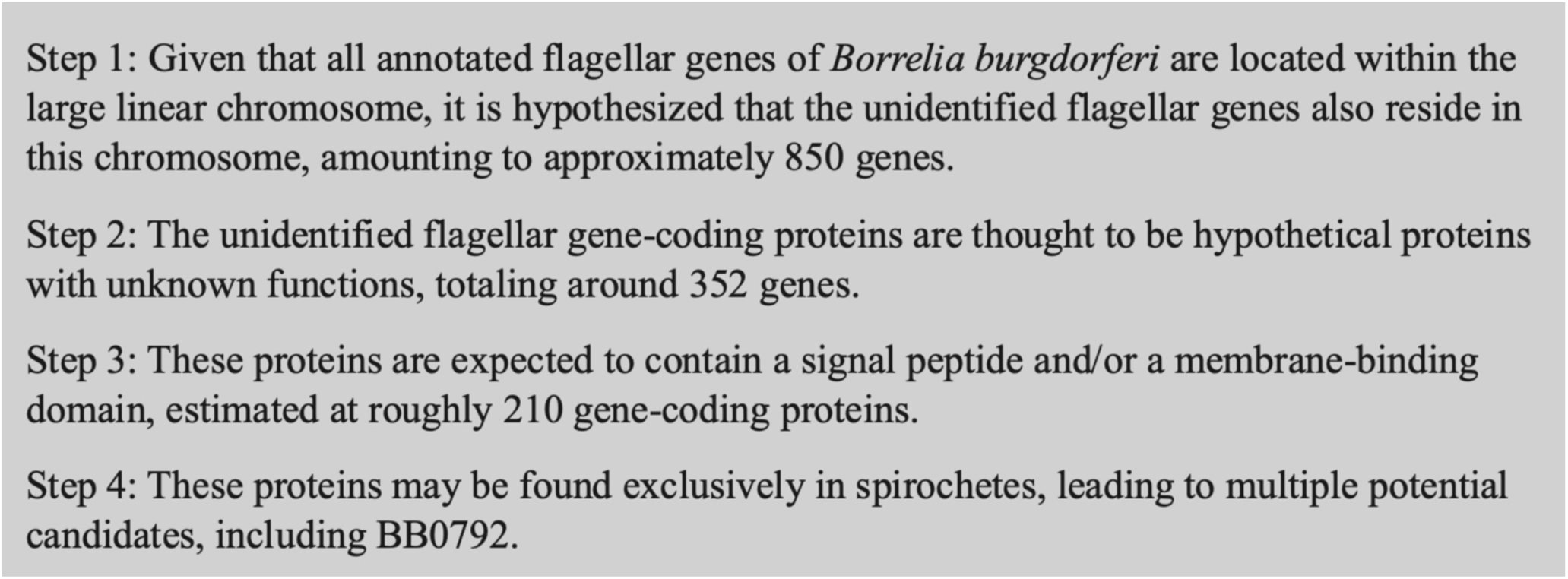
Candidate-screening strategy for identifying BB0792 as a putative flagellar component. The screening strategy began with approximately 850 genes on the large linear chromosome of *B. burgdorferi* and narrowed the candidate set to approximately 352 genes encoding hypothetical proteins. Further prioritization for proteins with a predicted signal peptide and/or membrane-binding domain yielded approximately 210 candidates. A final criterion of putative spirochete-specific distribution identified several candidates, including BB0792, subsequently designated FrbA.

**Extended Data Fig. 5:**
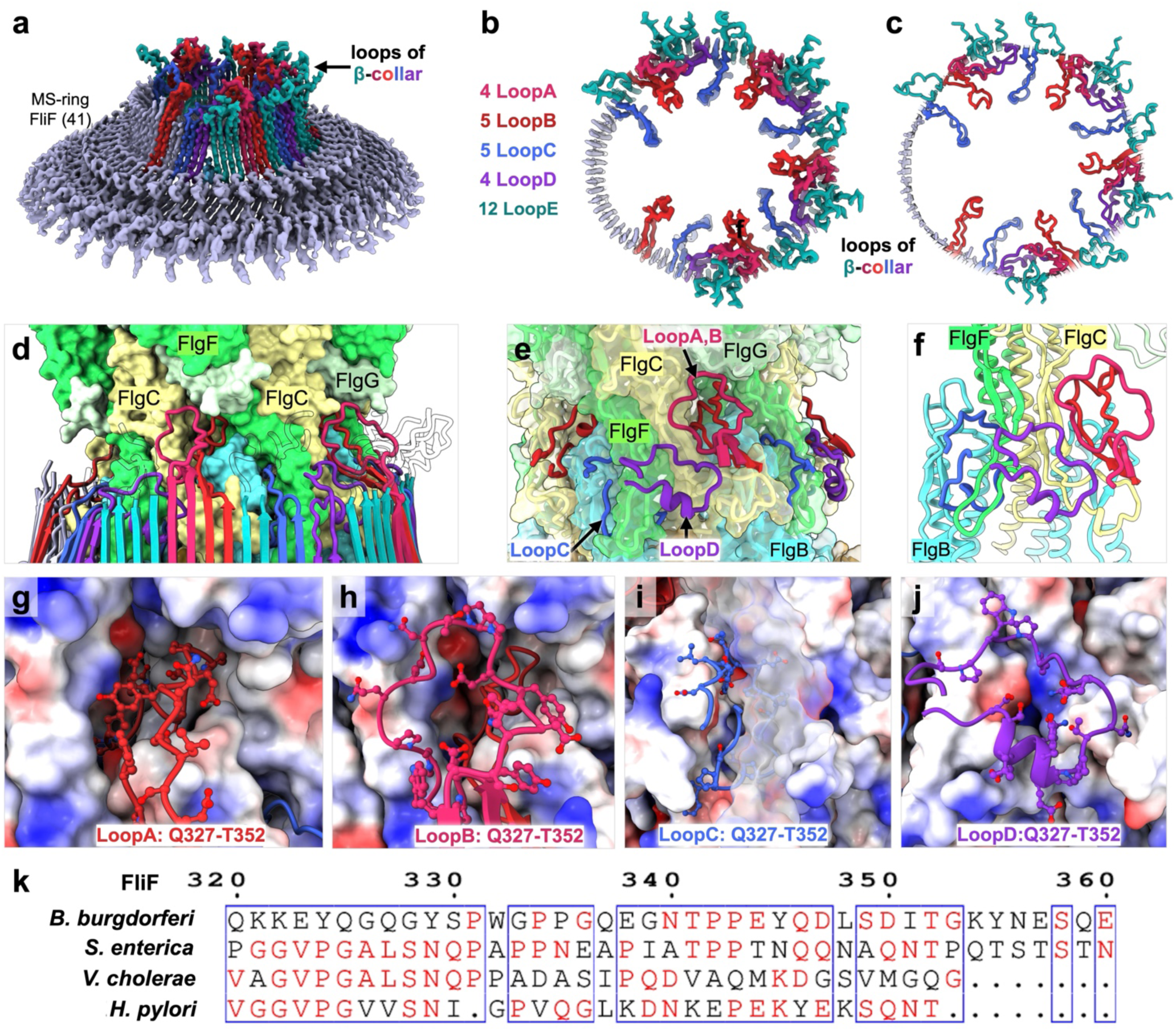
Conformational diversity of the FliF β-collar loops at the MS-ring–rod interface. **(a**) MS-ring density map highlighting distinct conformations of the FliF β-collar loops. **b**, **c**, Axial views of the loop densities (**b**) and corresponding atomic models (**c**). LoopA–LoopE are coloured red, pink, blue, purple and teal, respectively; the remainder of the MS-ring is light purple. **d**–**f**, Close-up views of the MS-ring–rod interface showing the FliF loops extending towards the rod in surface and ribbon representations. **g**–**j**, Detailed views of LoopA (**g**), LoopB (**h**), LoopD (**i**) and LoopC (**j**) at the rod surface. Loop side chains are shown in ball-and-stick representation, and the rod surface is coloured by electrostatic potential, with red and blue indicating negative and positive potential, respectively. **k**, Sequence alignment of the FliF loop region from *B. burgdorferi*, *Salmonella enterica*, *Vibrio cholerae* and *Helicobacter pylori*.

**Extended Data Fig. 6:**
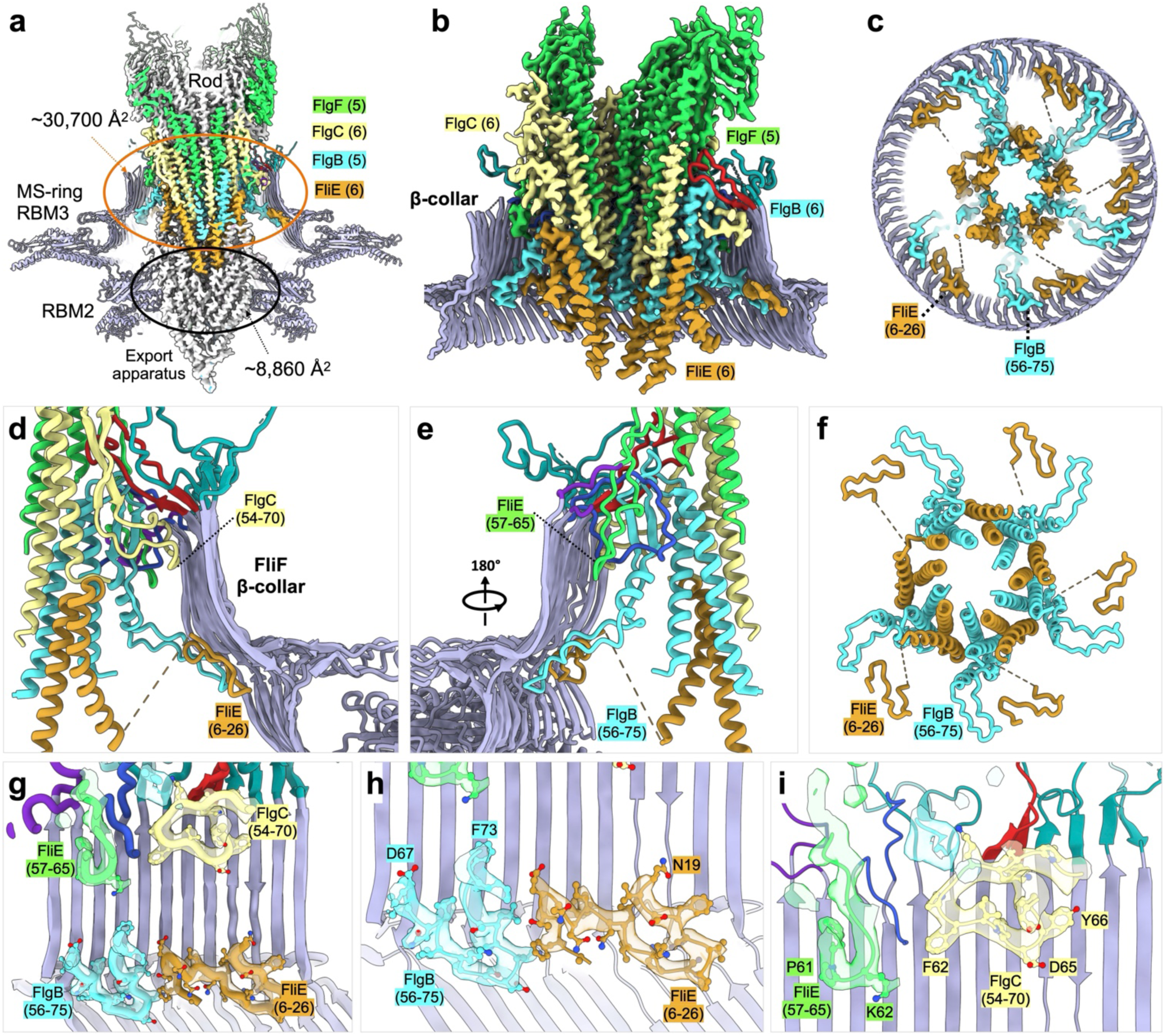
Structural contacts between proximal rod components and the FliF β-collar. **a**, Cutaway view of the MS-ring–rod density map showing the proximal rod within the MS-ring. Buried surface areas are indicated for the interfaces between the MS-ring loops and the rod, and between FliF RBM1/RBM2 and the FliPQR–FlhB export apparatus. **b**, Enlarged view of the MS-ring–rod interface, highlighting rod elements that extend towards the FliF β-collar. **c**, Axial section showing the arrangement of FliE and FlgB within the surrounding MS-ring. **d**, **e**, Ribbon views from different sides showing contacts between the rod and the inner surface of the β-collar. **f**, Axial view of the FliE–FlgB assembly. **g**, Overview of rod elements contacting the β-collar. **h**–**j**, Enlarged views highlighting contacts involving FliE and FlgB near the base of the β-collar (**h**), FlgF (green; **i**) and FlgC (pale yellow; **j**). RBM, ring-building motif.

**Extended Data Fig. 7:**
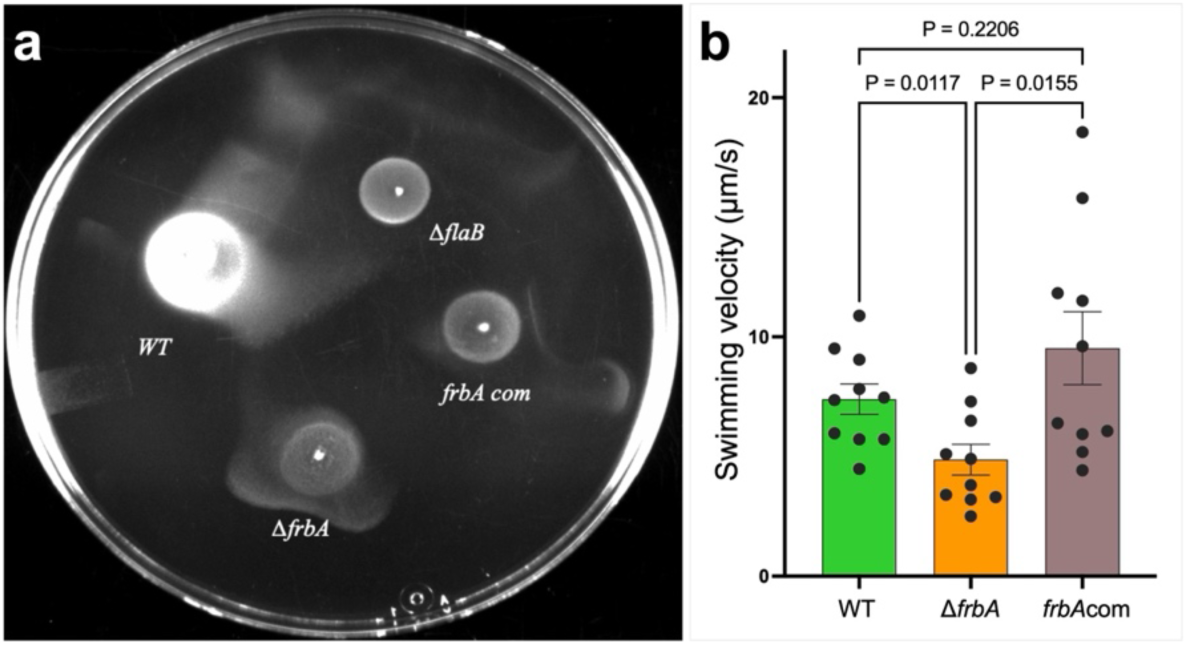
FrbA contributes to motility in *B. burgdorferi*. **a**, Representative swarm plate showing the motility of wild-type (WT), Δ*flaB*, Δ*frbA* and frbA-complemented (*frbA*com) strains. **b**, Swimming velocities of WT, Δ*frbA* and *frbA*com cells in in 0.5% methylcellulose. Data are mean ± s.e.; 10 cells were analyzed per strain. P values for the indicated comparisons were determined using Welch’s t-tests.

**Extended Data Fig. 8:**
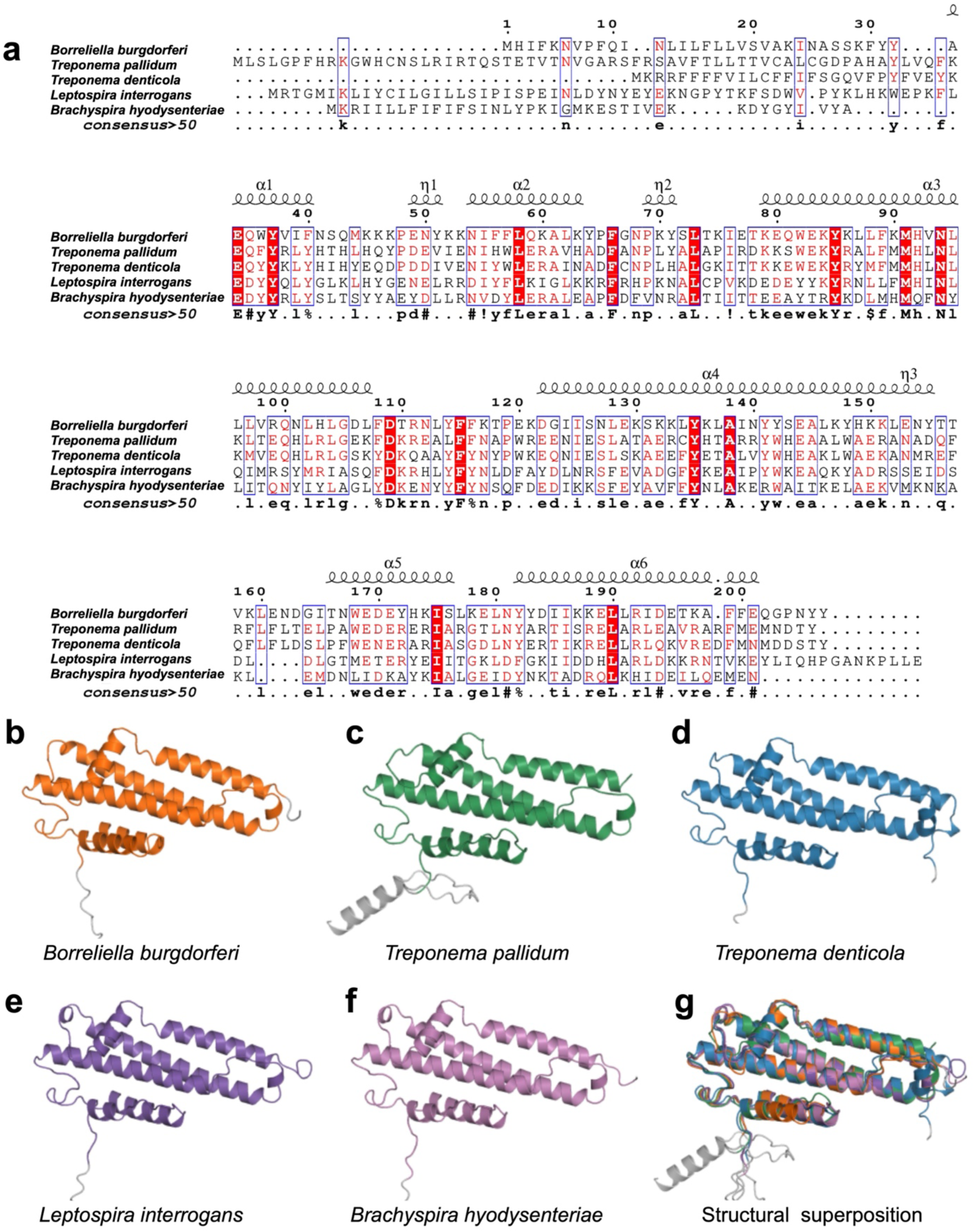
Sequence and structural conservation of FrbA across spirochetes. **a**, Multiple sequence alignment of FrbA homologues from *B. burgdorferi*, *Treponema pallidum*, *Treponema denticola*, *Leptospira interrogans* and *Brachyspira hyodysenteriae*. **b**–**f**, Ribbon representations of FrbA AlphaFold3 prediction structures from *B. burgdorferi* (**b**), *T. pallidum* (**c**), *T. denticola* (**d**), *L. interrogans* (**e**) and *B. hyodysenteriae* (**f**), shown in similar orientations. **g**, Superposition of the five models, colored as in **b**–**f**, showing a similar overall arrangement of the α-helical core.

**Extended Data Table 1.**
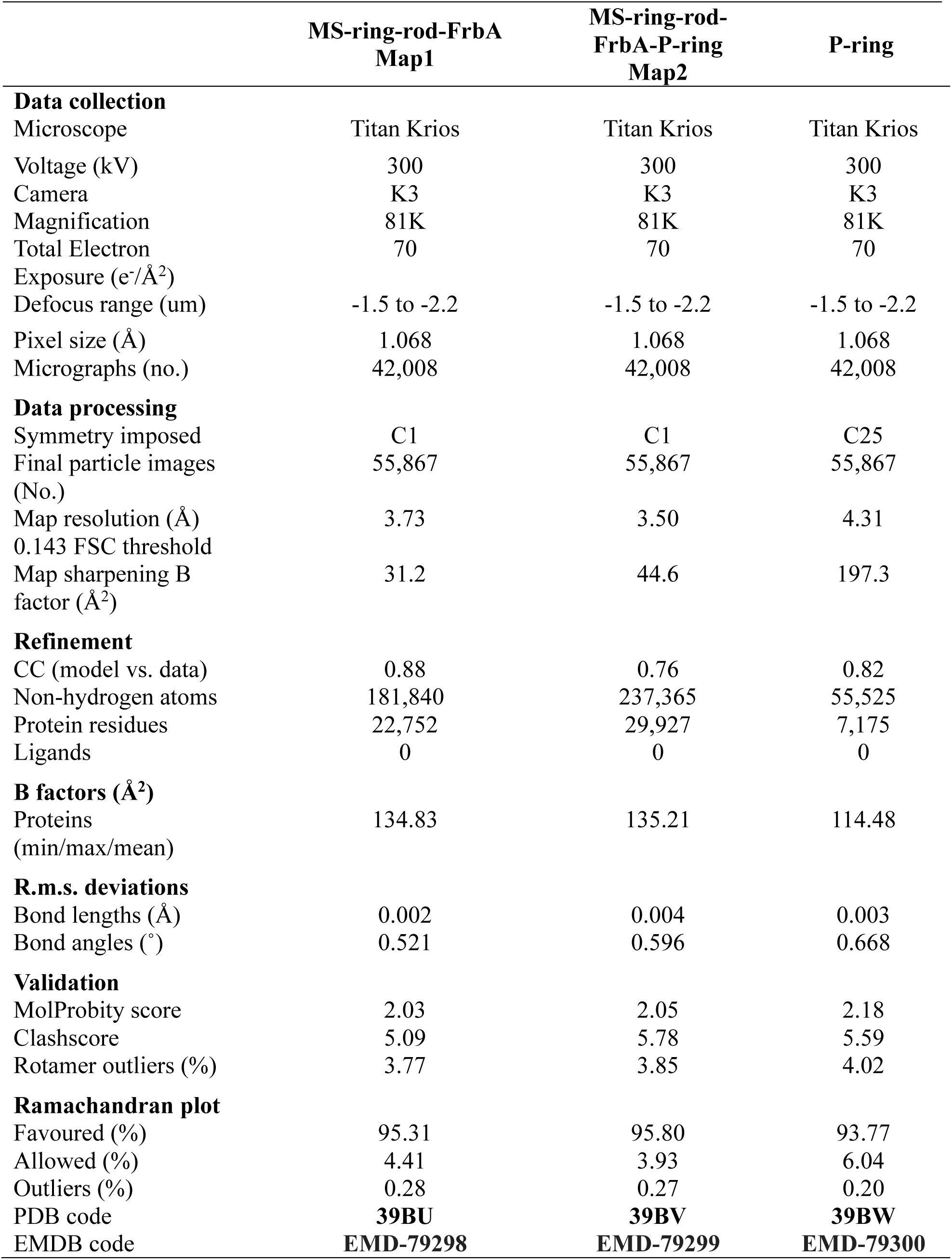
Data Collection and Structure Refinement Statistics.

## Reference

1 Wadhwa, N. & Berg, H. C. Bacterial motility: machinery and mechanisms. Nat Rev Microbiol 20, 161–173 (2022). 10.1038/s41579-021-00626-4

2 Hu, H. et al. Structural basis of torque generation in the bi-directional bacterial flagellar motor. Trends Biochem Sci 47, 160–172 (2022). 10.1016/j.tibs.2021.06.005

3 Guo, S. & Liu, J. The Bacterial Flagellar Motor: Insights Into Torque Generation, Rotational Switching, and Mechanosensing. Front Microbiol 13, 911114 (2022). 10.3389/fmicb.2022.911114

4 Chen, S. et al. Structural diversity of bacterial flagellar motors. EMBO J 30, 2972–2981 (2011). 10.1038/emboj.2011.186

5 Beeby, M. et al. Diverse high-torque bacterial flagellar motors assemble wider stator rings using a conserved protein scaffold. Proc Natl Acad Sci U S A 113, E1917–1926 (2016). 10.1073/pnas.1518952113

6 Chang, Y. et al. Molecular mechanism for rotational switching of the bacterial flagellar motor. Nat Struct Mol Biol 27, 1041–1047 (2020). 10.1038/s41594-020-0497-2

7 Drobnic, T. et al. In situ structure of a bacterial flagellar motor at subnanometre resolution reveals adaptations for increased torque. Nat Microbiol 10, 1723–1740 (2025). 10.1038/s41564-025-02012-9

8 Tachiyama, S. et al. The flagellar motor protein FliL forms a scaffold of circumferentially positioned rings required for stator activation. Proc Natl Acad Sci U S A 119 (2022). 10.1073/pnas.2118401119

9 Nakamura, S., Leshansky, A., Magariyama, Y., Namba, K. & Kudo, S. Direct measurement of helical cell motion of the spirochete *leptospira*. Biophys J 106, 47–54 (2014). 10.1016/j.bpj.2013.11.1118

10 Liu, J. et al. Intact flagellar motor of *Borrelia burgdorferi* revealed by cryo-electron tomography: evidence for stator ring curvature and rotor/C-ring assembly flexion. J Bacteriol 191, 5026–5036 (2009). 10.1128/JB.00340-09

11 Charon, N. W. et al. The flat-ribbon configuration of the periplasmic flagella of *Borrelia burgdorferi* and its relationship to motility and morphology. J Bacteriol 191, 600–607 (2009). 10.1128/JB.01288-08

12 Motaleb, M. A. et al. Borrelia burgdorferi periplasmic flagella have both skeletal and motility functions. Proc Natl Acad Sci U S A 97, 10899–10904 (2000). 10.1073/pnas.200221797

13 Charon, N. W. et al. The unique paradigm of spirochete motility and chemotaxis. Annu Rev Microbiol 66, 349–370 (2012). 10.1146/annurev-micro-092611-150145

14 Botting, J. M. et al. FlbB forms a distinctive ring essential for periplasmic flagellar assembly and motility in *Borrelia burgdorferi*. PLoS Pathog 21, e1012812 (2025). 10.1371/journal.ppat.1012812

15 Yue, J. et al. In situ structure of the flagellar export apparatus reveals membrane remodeling and lipid-mediated coupling between FlhA and the export gate. Nature Communications (2026). 10.1038/s41467-026-77713-8

16 Tan, J. et al. Structural basis of assembly and torque transmission of the bacterial flagellar motor. Cell 184, 2665–2679 e2619 (2021). 10.1016/j.cell.2021.03.057

17 Johnson, S. et al. Molecular structure of the intact bacterial flagellar basal body. Nat Microbiol 6, 712–721 (2021). 10.1038/s41564-021-00895-y

18 Zhang, L. et al. Molecular basis of high-torque transmission of the *Vibrio* polar flagellar motor. Protein Cell (2026). 10.1093/procel/pwag025

19 Zhao, X. et al. Cryoelectron tomography reveals the sequential assembly of bacterial flagella in *Borrelia burgdorferi*. Proc Natl Acad Sci U S A 110, 14390–14395 (2013). 10.1073/pnas.1308306110

20 Jamali, K. et al. Automated model building and protein identification in cryo-EM maps. Nature 628, 450–457 (2024). 10.1038/s41586-024-07215-4

21 Li, T., Chen, J., Li, H., Cao, H. & Huang, S. Y. EMProt improves structure determination from cryo-EM maps. Nat Struct Mol Biol 33, 341–350 (2026). 10.1038/s41594-025-01723-1

22 Abramson, J. et al. Accurate structure prediction of biomolecular interactions with AlphaFold 3. Nature 630, 493–500 (2024). 10.1038/s41586-024-07487-w

23 Oshiro, R. T., Rajendren, S., Hundley, H. A. & Kearns, D. B. Robust Stoichiometry of FliW-CsrA Governs Flagellin Homeostasis and Cytoplasmic Organization in Bacillus subtilis. mBio 10 (2019). 10.1128/mBio.00533-19

24 Thomson, N. M. & Pallen, M. J. Restoration of wild-type motility to flagellin-knockout *Escherichia coli* by varying promoter, copy number and induction strength in plasmid-based expression of flagellin. Curr Res Biotechnol 2, 45–52 (2020). 10.1016/j.crbiot.2020.03.001

25 Delalez, N. J., Berry, R. M. & Armitage, J. P. Stoichiometry and turnover of the bacterial flagellar switch protein FliN. mBio 5, e01216–01214 (2014). 10.1128/mBio.01216-14

26 Tachiyama, S. et al. FlgY, PflA, and PflB form a spoke-ring network in the high-torque flagellar motor of *Helicobacter pylori*. Proc Natl Acad Sci U S A 122, e2421632122 (2025). 10.1073/pnas.2421632122

27 Harman, M. W. et al. The heterogeneous motility of the Lyme disease spirochete in gelatin mimics dissemination through tissue. Proc Natl Acad Sci U S A 109, 3059–3064 (2012). 10.1073/pnas.1114362109

28 Picardeau, M. Virulence of the zoonotic agent of leptospirosis: still terra incognita? Nat Rev Microbiol 15, 297–307 (2017). 10.1038/nrmicro.2017.5

29 Nakamura, S. Spirochete Flagella and Motility. Biomolecules 10 (2020). 10.3390/biom10040550

30 Sultan, S. Z., Pitzer, J. E., Miller, M. R. & Motaleb, M. A. Analysis of a *Borrelia burgdorferi* phosphodiesterase demonstrates a role for cyclic-di-guanosine monophosphate in motility and virulence. Mol Microbiol 77, 128–142 (2010). 10.1111/j.1365-2958.2010.07191.x

31 Motaleb, M. A., Miller, M. R., Bakker, R. G., Li, C. & Charon, N. W. Isolation and characterization of chemotaxis mutants of the Lyme disease Spirochete *Borrelia burgdorferi* using allelic exchange mutagenesis, flow cytometry, and cell tracking. Methods Enzymol 422, 421–437 (2007). 10.1016/S0076-6879(06)22021-4

32 Li, X. et al. The Lyme disease agent *Borrelia burgdorferi* requires BB0690, a Dps homologue, to persist within ticks. Mol Microbiol 63, 694–710 (2007). 10.1111/j.1365-2958.2006.05550.x

33 Pitzer, J. E. et al. Analysis of the *Borrelia burgdorferi* cyclic-di-GMP-binding protein PlzA reveals a role in motility and virulence. Infect Immun 79, 1815–1825 (2011). 10.1128/IAI.00075-11

34 Zhang, X., Yang, X., Kumar, M. & Pal, U. BB0323 function is essential for *Borrelia burgdorferi* virulence and persistence through tick-rodent transmission cycle. J Infect Dis 200, 1318–1330 (2009). 10.1086/605846

35 Yang, X. et al. Characterization of unique regions of *Borrelia burgdorferi* surface-located membrane protein 1. Infect Immun 78, 4477–4487 (2010). 10.1128/IAI.00501-10

36 Yang, X., Coleman, A. S., Anguita, J. & Pal, U. A chromosomally encoded virulence factor protects the Lyme disease pathogen against host-adaptive immunity. PLoS Pathog 5, e1000326 (2009). 10.1371/journal.ppat.1000326

37 Sultan, S. Z. et al. Motility is crucial for the infectious life cycle of *Borrelia burgdorferi*. Infect Immun 81, 2012–2021 (2013). 10.1128/IAI.01228-12

38 Sultan, S. Z. et al. Motor rotation is essential for the formation of the periplasmic flagellar ribbon, cellular morphology, and *Borrelia burgdorferi* persistence within Ixodes scapularis tick and murine hosts. Infect Immun 83, 1765–1777 (2015). 10.1128/IAI.03097-14

39 Motaleb, M. A., Pitzer, J. E., Sultan, S. Z. & Liu, J. A novel gene inactivation system reveals altered periplasmic flagellar orientation in a *Borrelia burgdorferi* fliL mutant. J Bacteriol 193, 3324–3331 (2011). 10.1128/JB.00202-11

40 Guo, W. et al. Near-atomic in situ architecture and membrane-coupled dynamics of the *Vibrio cholerae* sheathed flagellum. Nat Commun 17 (2026). 10.1038/s41467-026-75593-6

41 Yue, J. et al. In situ structures of the Legionella Dot/Icm T4SS identify the DotA-IcmX complex as the gatekeeper for effector translocation. Proc Natl Acad Sci U S A 122, e2516300122 (2025). 10.1073/pnas.2516300122

42 Mastronarde, D. N. Automated electron microscope tomography using robust prediction of specimen movements. J Struct Biol 152, 36–51 (2005). 10.1016/j.jsb.2005.07.007

43 Punjani, A., Rubinstein, J. L., Fleet, D. J. & Brubaker, M. A. cryoSPARC: algorithms for rapid unsupervised cryo-EM structure determination. Nat Methods 14, 290–296 (2017). 10.1038/nmeth.4169

44 Bepler, T. et al. Positive-unlabeled convolutional neural networks for particle picking in cryo-electron micrographs. Nat Methods 16, 1153–1160 (2019). 10.1038/s41592-019-0575-8

45 Cao, H. et al. EMReady2: improvement of cryo-EM and cryo-ET maps by local quality-aware deep learning with Mamba. Nature Communications 17, 6493 (2026). 10.1038/s41467-026-71794-1

46 Burt, A. et al. An image processing pipeline for electron cryo-tomography in RELION-5. FEBS Open Bio 14, 1788–1804 (2024). 10.1002/2211-5463.13873

47 Goddard, T. D. et al. UCSF ChimeraX: Meeting modern challenges in visualization and analysis. Protein Sci 27, 14–25 (2018). 10.1002/pro.3235

48 Kidmose, R. T. et al. Namdinator - automatic molecular dynamics flexible fitting of structural models into cryo-EM and crystallography experimental maps. IUCrJ 6, 526–531 (2019). 10.1107/S2052252519007619

49 Croll, T. I. ISOLDE: a physically realistic environment for model building into low-resolution electron-density maps. Acta Crystallogr D Struct Biol 74, 519–530 (2018). 10.1107/S2059798318002425

50 Emsley, P. & Cowtan, K. Coot: model-building tools for molecular graphics. Acta Crystallogr D Biol Crystallogr 60, 2126–2132 (2004). 10.1107/S0907444904019158

51 Liebschner, D. et al. Macromolecular structure determination using X-rays, neutrons and electrons: recent developments in Phenix. Acta Crystallogr D Struct Biol 75, 861–877 (2019). 10.1107/S2059798319011471

52 Davis, I. W. et al. MolProbity: all-atom contacts and structure validation for proteins and nucleic acids. Nucleic Acids Res 35, W375–383 (2007). 10.1093/nar/gkm216

53 Xu, A. & Xu, C. FastTomo: A SerialEM Script for Collecting Electron Tomography Data. bioRxiv (2021). 10.1101/2021.03.16.435675

54 Zheng, S. Q. et al. MotionCor2: anisotropic correction of beam-induced motion for improved cryo-electron microscopy. Nat Methods 14, 331–332 (2017). 10.1038/nmeth.4193

55 Kremer, J. R., Mastronarde, D. N. & McIntosh, J. R. Computer visualization of three-dimensional image data using IMOD. J Struct Biol 116, 71–76 (1996). 10.1006/jsbi.1996.0013

56 Zhang, K. Gctf: Real-time CTF determination and correction. J Struct Biol 193, 1–12 (2016). 10.1016/j.jsb.2015.11.003

57 Agulleiro, J. I. & Fernandez, J. J. Tomo3D 2.0--exploitation of advanced vector extensions (AVX) for 3D reconstruction. J Struct Biol 189, 147–152 (2015). 10.1016/j.jsb.2014.11.009

58 Winkler, H. et al. Tomographic subvolume alignment and subvolume classification applied to myosin V and SIV envelope spikes. Journal of Structural Biology 165, 64–77 (2009). 10.1016/j.jsb.2008.10.004

59 Feng, X. et al. Structural insights into the assembly and evolution of a complex bacterial flagellar motor. Nat Microbiol 11, 770–785 (2026). 10.1038/s41564-025-02248-5

60 Rajkovic, A. et al. HMMER web server: 2026 update. Nucleic Acids Res 54, W272–W278 (2026). 10.1093/nar/gkag373

61 van Kempen, M. et al. Fast and accurate protein structure search with Foldseek. Nat Biotechnol 42, 243–246 (2024). 10.1038/s41587-023-01773-0

62 Pettersen, E. F. et al. UCSF ChimeraX: Structure visualization for researchers, educators, and developers. Protein Sci 30, 70–82 (2021). 10.1002/pro.3943

63 Parks, D. H. et al. GTDB release 10: a complete and systematic taxonomy for 715 230 bacterial and 17 245 archaeal genomes. Nucleic Acids Res 54, D743–D754 (2026). 10.1093/nar/gkaf1040

64 Letunic, I. & Bork, P. Interactive Tree of Life (iTOL) v6: recent updates to the phylogenetic tree display and annotation tool. Nucleic Acids Res 52, W78–W82 (2024). 10.1093/nar/gkae268

